# Inferring fitness seascapes from evolutionary histories

**DOI:** 10.1101/2025.06.08.658500

**Authors:** Yirui Gao, Brian Lee, John P. Barton

## Abstract

Evolutionary dynamics are often conceptualized as walks on an adaptive landscape, with populations climbing toward optimal fitness peaks. However, environmental changes can transform static landscapes into dynamic “fitness seascapes” where natural selection fluctuates in time. Here, we used a path integral approach derived from statistical physics to reveal time-varying selection pressures from genetic sequence data. We found that constraints on the fitness seascape are determined by the screened Poisson equation, which also describes the screening of electric fields in media with mobile charge carriers. In our model, changes in mutation frequencies act as “charges” that reveal the underlying fitness seascape, which is analogous to an electrostatic potential. After validating our method in simulations, we applied it to study how HIV-1 evolves to escape immune control by T cells within individual hosts. Our analysis showed that the fitness benefit of immune escape declines as T cell responses approach their peak intensity, suggesting that functional exhaustion may impair the effectiveness of the immune response against HIV-1. Overall, our approach provides a general framework for capturing the complex dynamics of natural selection in rapidly evolving populations.

## Introduction

The concept of natural selection is central to evolution. Reproductively successful individuals pass on their genes to subsequent generations, driving genetic change through the “survival of the fittest.” This process is often visualized through fitness landscapes, where each point represents a possible genotype and its elevation corresponds to reproductive fitness. In this metaphor, evolution can be seen as populations climbing uphill toward fitness peaks while avoiding low-fitness valleys.

However, the fitness effects of traits or mutations can change along with the environment. In one of the most famous examples of natural selection, darkly colored moths became favored over lightly pigmented variants during the Industrial Revolution, as their colors provided better camouflage in smoky environments ^1,2^. When pollution levels declined, lightly colored moths returned. Modern analyses have also highlighted fluctuating selection in other natural populations ^3–8^. This dynamic is captured by the concept of fitness seascapes, where the landscape itself shifts over time ^9^.

Temporal genetic data – sequences sampled over time from sources such as experimental evolution, genomic surveillance of pathogens, or ancient DNA ^10–13^– provide valuable windows into population dynamics that can be leveraged to understand evolution. These data capture changes in the genetic structure of populations as they occur in response to shifting fitness constraints. However, signals of adaptation must be carefully separated from other stochastic contributions to population dynamics.

At present, several methods have been developed to learn fluctuating fitness effects from data ^8,14–16^. One limitation of current approaches is that they focus on the fitness effects of single mutations, without explicitly considering the effects of the genetic background. This simplification risks overlooking important contributions to evolution. For example, beneficial mutations can be driven to extinction through competition with even fitter variants in a process known as clonal interference ^17^. Conversely, neutral mutations can hitchhike to high frequencies when they appear on high-fitness genetic backgrounds ^18^. These complex interactions highlight the need for approaches that account for the genetic context in which evolution occurs. Alternatively, a parallel line of work has studied how fitness (and changes in fitness) is reflected in evolutionary dynamics over time, but without estimating the individual contributions of specific mutations ^19,20^.

Here, we developed a solution to this problem using methods from statistical physics. Starting from a stochastic model of population evolution, we derived a path integral that quantifies the relative likelihood of different evolutionary histories. One can then obtain an analytical equation for the time-varying fitness effects of mutations that best explain the observed trajectory of evolution. Remarkably, this expression maps onto the screened Poisson equation from physics. In our inference framework, time plays the role of a spatial coordinate, and the fitness effects of mutations are like an electric potential. Changes in the frequencies of mutations act like charges that describe the shape of the fitness seascape at a particular moment in time. We validate our approach in simulations and apply it to study how human immunodeficiency virus (HIV)-1 evolves within individual hosts.

## Results

### Evolutionary model

Our starting point for analysis is the classic Wright-Fisher (WF) model ^21^, which describes the dynamics of a population of *N* individuals subject to mutations, genetic recombination, and natural selection. For simplicity, we will represent the haploid genetic sequence of each individual *a* as a binary string 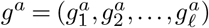 with 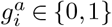. Here the index *i* labels each site in the genetic sequence, running from 1 to 𝓁, and *a* labels the genotype (of which there are *M* = 2^𝓁^ possibilities). The probability of mutation per site per reproduction cycle is *µ*, which we fix in time. Mutation at a site *i* converts the value of the genetic sequence at that site, *g*_*i*_, from the wild-type (WT, represented by 0) to the mutant (1) state, or vice versa.

Recombination refers to the process of genetic exchange between individuals, producing a “child” genome with shuffled contributions from each of the “parents.” While this phenomenon is most familiar in sexual reproduction, it can also occur in other contexts, including HIV-1 replication within host cells ^22^. We write the probability of a recombination breakpoint occurring between any adjacent sites in the genome as *r*, which we also assume to be time-independent. When recombination occurs between two parents *a* and *b* at site *i*, the resulting offspring *c* inherits positions 1 through *i* from parent *a* and *i* + 1 through 𝓁 from parent *b*, creating the sequence 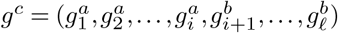.

To model natural selection, we assign a time-dependent fitness value *f*_*a*_(*t*) to each genotype *a*. This fitness value directly influences reproductive success: genotypes with higher fitness values contribute more offspring to the next generation. We can express the fitness function as a polynomial expansion of genetic contributions,

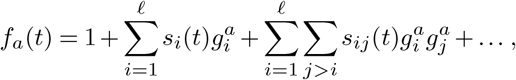

where the first-order terms *s*_*i*_(*t*) represent the direct fitness effects of individual mutations and higher-order terms capture interactions between mutations. While this framework can accommodate arbitrarily complex fitness landscapes with many interaction terms, we focus on the simpler case where only first-order terms *s*_*i*_(*t*) are included. This approximation captures the essential dynamics of a shifting fitness seascape while remaining mathematically tractable. In population genetics, the *s*_*i*_(*t*) are called selection coefficients and measure the fitness advantage or disadvantage of each mutation.

We define *n*_*a*_(*t*) as the number of individuals with genotype *a* at time *t*, and *z*_*a*_(*t*) = *n*_*a*_(*t*)*/N* as the corresponding frequency. The state of the population is described by a genotype frequency vector ***z***(*t*) = (*z*_1_(*t*), *z*_2_(*t*),…, *z*_*M*_ (*t*)). In the WF model, evolution proceeds stochastically. The state of the population in the next generation, ***z***(*t* + 1), follows a multinomial distribution

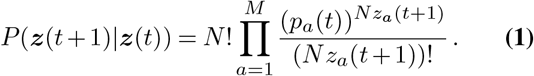

Here, *p*_*a*_(*t*) represents the expected frequency of genotype *a* in the next generation, incorporating the effects of natural selection, mutation, and recombination (with detailed expressions presented in Supplementary Information).

### Inference of fitness seascapes

In principle, we could use the transition probability in equation (1) to estimate selection coefficients from observed evolutionary trajectories. Following principles of Bayesian inference ^23^, the posterior probability of the selection coefficients ***s***(*t*) = (*s*_1_(*t*), *s*_2_(*t*),…, *s*_𝓁_(*t*)) from one generation of evolution is

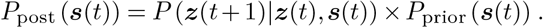

The first term represents how likely we are to observe the actual evolutionary trajectory given particular selection coefficients, and *P*_prior_ (***s***(*t*)) encodes our prior knowledge about plausible selection coefficient values. However, the functional form of (1) is complicated, making this expression difficult to work with directly.

To make this problem tractable, we apply the diffusion approximation of the WF model ^21,24,25^. This mathematical approach considers the scaling limit in which the population size *N* → ∞, while the selection coefficients, mutation rate, and recombination rate are all 𝒪(1*/N*) (Supplementary Information; see also ref. ^26^). In this limit, we obtain a continuous-time stochastic model of population evolution. The probability of an evolutionary trajectory, or “path,” of genotype frequencies from times *t*_0_ to *t*_*K*_, conditioned on the initial state of the population ***z***(*t*_0_), can then be expressed as a path integral ^26^ (see also refs. ^27,28^)

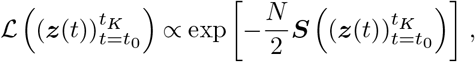

with

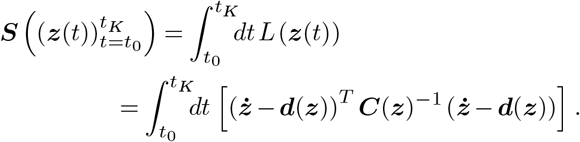

Here and below, we suppress the time indices on the genotype frequency vectors ***z***(*t*) and other time-varying quantities when their time dependence is clear. In the above expression, ***d***(***z***) is the drift vector, representing the deterministic forces of selection, mutation, and recombination driving evolution, and ***C***(***z***) is the diffusion matrix, which captures stochastic fluctuations (Supplementary Information).

Imposing additional constraints on the ***s***(*t*) through the prior distribution *P*_prior_ (***s***(*t*)) can improve our ability to estimate realistic selection coefficients. We assume that most mutations have little effect on fitness, which we quantify through a Gaussian distribution with mean zero and variance 1*/Nγ* for the selection coefficients,

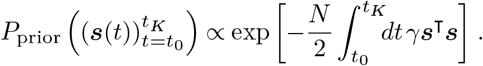

Larger values of *γ* penalize large selection coefficients more heavily, shrinking estimates toward zero unless they are well-supported by data.

Second, we introduce another biologically realistic constraint: environmental changes (and consequently selection pressures) typically do not fluctuate arbitrarily quickly. We implement this through a Gaussian prior distribution for the time derivative of the selection coefficients with mean zero and variance 1*/Nγ*′,

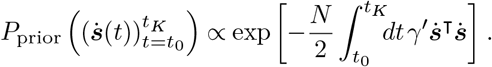

This term penalizes abrupt changes in the ***s***(*t*), with *γ*′ determining the expected smoothness in time. Larger values of *γ*′ are consistent with a gradually changing environment, while smaller values allow for more rapid fluctuations.

Combining the likelihood of the evolutionary dynamics and the prior constraints, we can derive an expression for the selection coefficients ŝ(*t*) that best explain an observed evolutionary trajectory. This maximum *a posteriori* (MAP) estimate is given by the solution of the Euler-Lagrange equation (Supplementary Information)

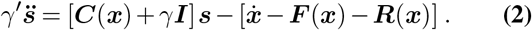

The vector ***x***(*t*) = (*x*_1_(*t*), *x*_2_(*t*),…, *x*_𝓁_(*t*)) represents the mutant frequency at each site in the genome across the population at time *t*,

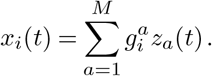

In (2), ***I*** is the identity matrix and ***C***(***x***(*t*)) is the mutant frequency covariance matrix, with entries

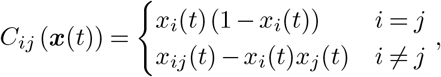

where 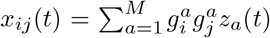. In population genetics, the covariance between mutant frequencies is referred to as linkage disequilibrium. Finally, ***F*** (***x***(*t*)) and ***R***(***x***(*t*)) quantify instantaneous change in mutant frequencies due to spontaneous mutations and recombination, respectively. In the model described above, *F*_*i*_(***x***(*t*)) = *µ* (1 − 2*x*_*i*_(*t*)) and ***R***(***x***(*t*)) = 0. The contribution from recombination is typically nonzero in more complex fitness models that include cooperative effects of mutations at multiple sites in the genome ^29,30^.

### Interpreting the Euler-Lagrange equation

Equation (2) provides an explicit expression linking evolutionary dynamics to evidence for a shifting fitness seascape. Under the diffusion approximation of the WF model, the expected instantaneous change in the mutant frequencies, 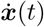, is given by (Supplementary Information)

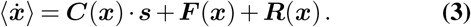

Thus, if the ***s***(*t*) are totally unconstrained (*γ* = *γ*′ = 0), the most likely selection coefficients would simply be those that exactly reproduce the observed changes mutant frequencies at each moment in time. However, this approach would yield noisy and biologically implausible results.

In reality, environmental changes and the resulting shifts in selection often occur gradually (captured by *γ*′ *>* 0). This implies that changes in mutant frequencies reveal information about selection not only at that time point, but also at times in the near past and future. To see this relationship more clearly, we can rearrange (2) as follows (**Fig. 1a**):

**Fig. 1.**
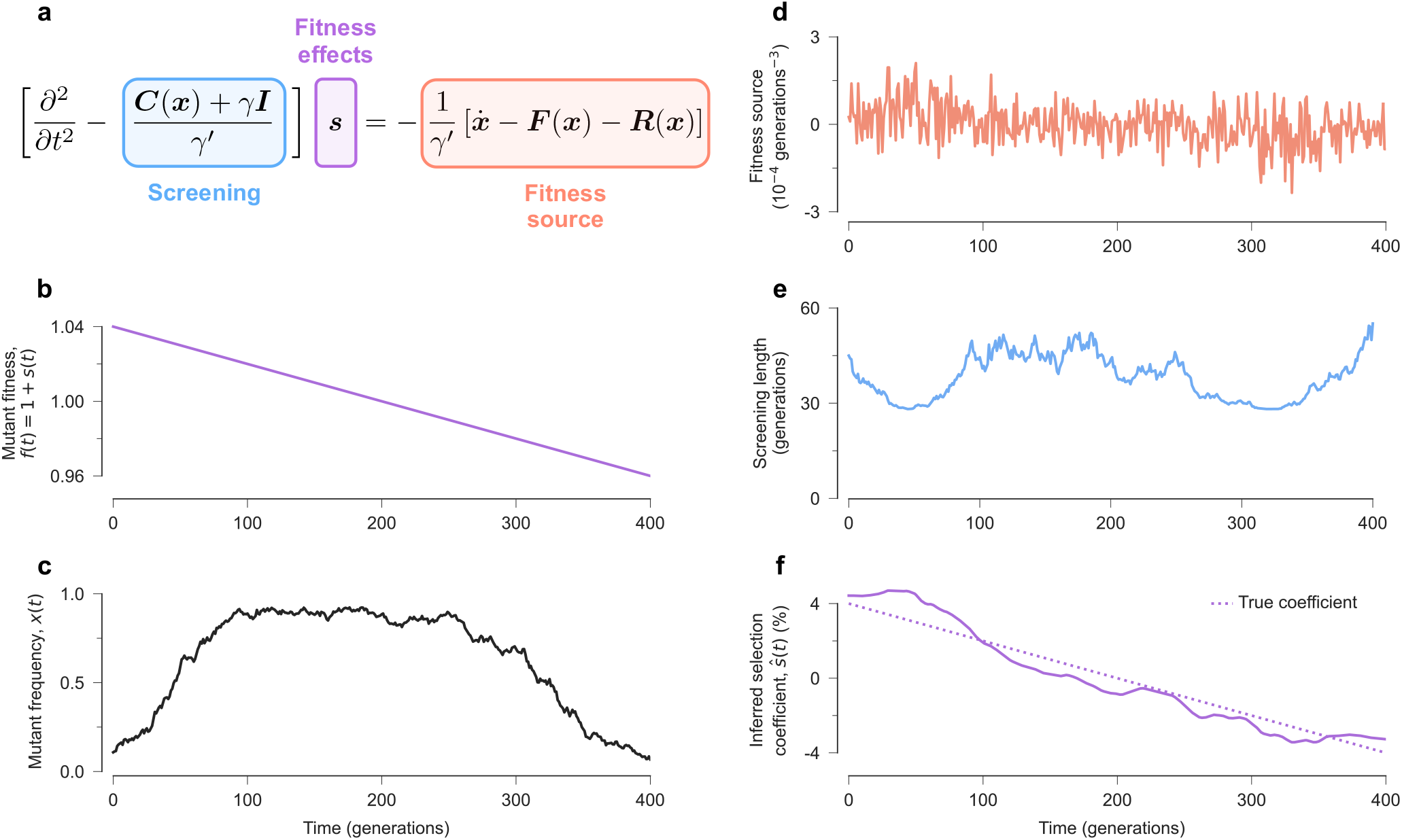
From temporal data to fitness seascapes. **a**, The time-varying fitness effects of mutations that best fit a particular data set, ***s*** (*t*), obey a screened Poisson equation. **b**, As an example, we consider a simple model with a single site (𝓁 = 1), where the fitness effect of mutations shifts from beneficial to deleterious over time. **c**, In an example simulation, mutant frequencies rise and then fall following changes in fitness. Instantaneous changes in mutant frequencies are a fitness “source” that provide evidence of natural selection (**d**), which propagate over a range of time determined by the screening length (**e**). **f**, Collectively, these terms guide our inference of natural selection over time, which closely follows the true, underlying seascape.

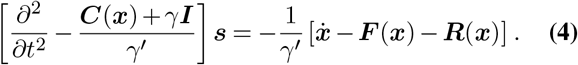

Interpreting time as a spatial coordinate, the Euler-Lagrange equation has the same form as the screened Poisson equation,

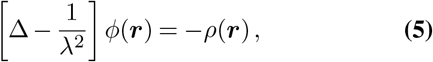

which arises in the context of electrostatic screening and granular flow ^31,32^, among other examples.

In the electrostatic context, (5) describes how electric fields are screened in a material with mobile charges. Here, *ϕ*(***r***) represents the electric potential, *ρ*(***r***) is the source term, and ***r*** are spatial coordinates. *λ* describes the characteristic distance over which screening occurs.

We can therefore provide an intuitive understanding of (4) by relating this expression with the screened Poisson equation in electrostatics (see **Fig. 1** for an example). In this analogy, the shape of the fitness seascape is defined by the selection coefficients in time, ***s***(*t*), which play the role of an electric potential in space, *ϕ*(***r***). The fitness seascape drives population evolution, as shown in (3). Changes in mutant frequency, 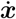, which are not explained by spontaneous mutations or recombination, ***F*** (***x***) + ***R***(***x***), are like charges that provide evidence of natural selection. Intuitively, beneficial mutations are likely to increase in frequency while deleterious ones decline. The screening length *λ* is proportional to 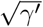, such that the effective time over which frequency changes are informative about fitness depends on how rapidly the environment changes.

The mutant frequency covariance matrix ***C***(***x***) plays multiple roles in (4). In the simplest case, one can imagine a diagonal covariance matrix, such that the mutations at each site in the genome are uncorrelated. In population genetics, this is referred to as linkage equilibrium. The mutant frequency variance *C*_*ii*_(*x*_*i*_(*t*)) is then inversely proportional to the screening length for *s*_*i*_. Thus, frequency changes near the boundaries (*x*_*i*_(*t*) near zero or one) propagate evidence of natural selection further in time.

The off-diagonal terms of ***C***(***x***) describe how correlations between mutations affect the response to natural selection. Expanding the first term on the right hand side of into diagonal and off-diagonal contributions, one can see that changes in mutant frequencies are driven not just by the fitness effect of the mutation, but also by the net fitness effects of the genetic background. For example, even if a mutation at some site *i* has no effect on fitness (*s*_*i*_(*t*) = 0), we would expect 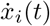 to be positive if mutations at site *i* were correlated with a beneficial mutation at another site *j* (*C*_*ij*_(***x***(*t*)) = *x*_*ij*_(*t*) − *x*_*i*_(*t*)*x*_*j*_(*t*) *>* 0 and *s*_*j*_ *>* 0).

### Treatment of boundary conditions

We applied the domain extension method to address boundary conditions in the screened Poisson equation. This approach involves expanding the original domain Ω to a larger extended domain 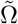, imposing new boundary conditions on 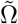, setting the source term to zero in 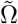, and using continuity conditions to connect solutions across domains. This approach is especially helpful for preventing artefacts at the edges of the observed time window. We extended the time domain and imposed a transversality condition 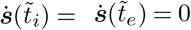, where 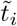 and 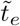 denote the initial and end times in the extended domain 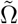, respectively (Supplementary Information). The transversality condition in our model is analogous to a Neumann boundary condition in electrostatics, which typically implies no flux through the boundary of an isolated or insulated domain. In our evolutionary context, this condition specifies that the environment is stationary at both the beginning and end of the extended time domain, rather than changing abruptly at the boundaries.

### Validation in simulations

To validate our method, we simulated population evolution on a simple, additive fitness seascape (**Fig. 2a**). Our model included *N* = 10^3^ individuals with genomes of length 𝓁 = 10. Mutations at six sites had fitness effects that were constant in time: two beneficial, two neutral, and two deleterious, with *s*(*t*) = 2%, 0, −2%, respectively. Mutations at four sites had time-varying effects on fitness: two each with selection coefficients of *s*(*t*) = *A* sin (2*πt/τ*) and *s*(*t*) = *A* cos (2*πt/τ*), with amplitude *A* = 4% and period *τ* = 1000 generations. Our simulations included mutations and recombination with rates *µ* = *r* = 1 × 10^−3^ per site per generation.

**Fig. 2.**
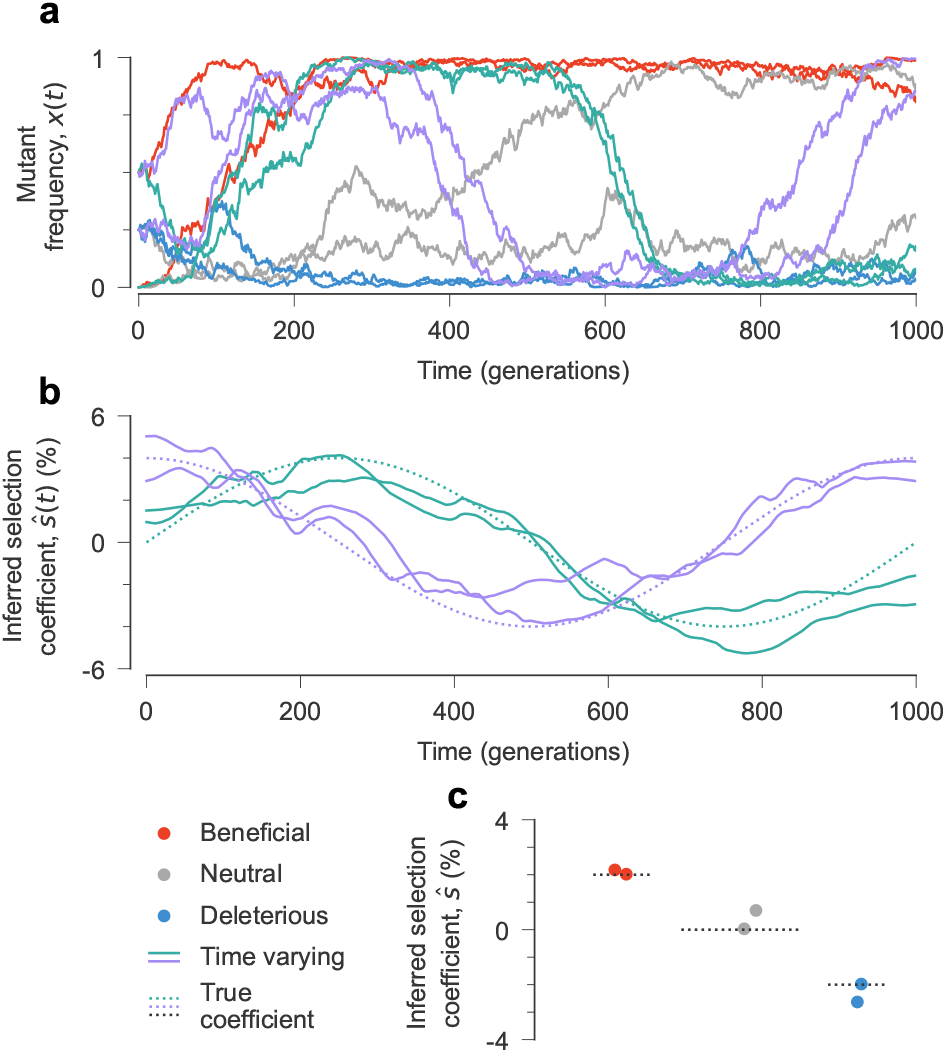
Inference of a simple fitness seascape. **a**, In simulated population dynamics, mutant frequencies change over time, driven by the fitness seascape. The inferred selection coefficients are close to the true, underlying ones for both time-varying parameters (**b**) and constant ones (**c**). Simulation parameters: 𝓁 = 10 sites with two states at each site (mutant and wild type, WT), 2 beneficial mutations with *s* = 0.02, 2 neutral mutations with *s* = 0, and 2 deleterious mutations with *s* = −0.02. Four selection coefficients follow a sine or cosine pattern over time. Population size *N* = 10^3^, mutation rate *µ* = 1 *×* 10^−3^ per site per generation, recombination probability *r* = 1 *×* 10^−3^ per site per generation. The initial population was randomly generated and evolved over *T* = 1000 generations.

**Figure 2b-c** displays results from a typical simulation, showing that our method can distinguish between beneficial, neutral, and deleterious mutations with constant effects and accurately estimate fluctuating selection. We performed 100 independent simulations to verify the consistency of our results (**Fig. 3**). We further tested the robustness of our approach to model misspecification, i.e., incorrectly assuming time-varying selection at sites where selection is actually constant. **Supplementary Fig. S1** shows that, even with this incorrect assumption, our method recovers selection coefficients that fluctuate around the true, constant values rather than producing spurious patterns.

**Fig. 3.**
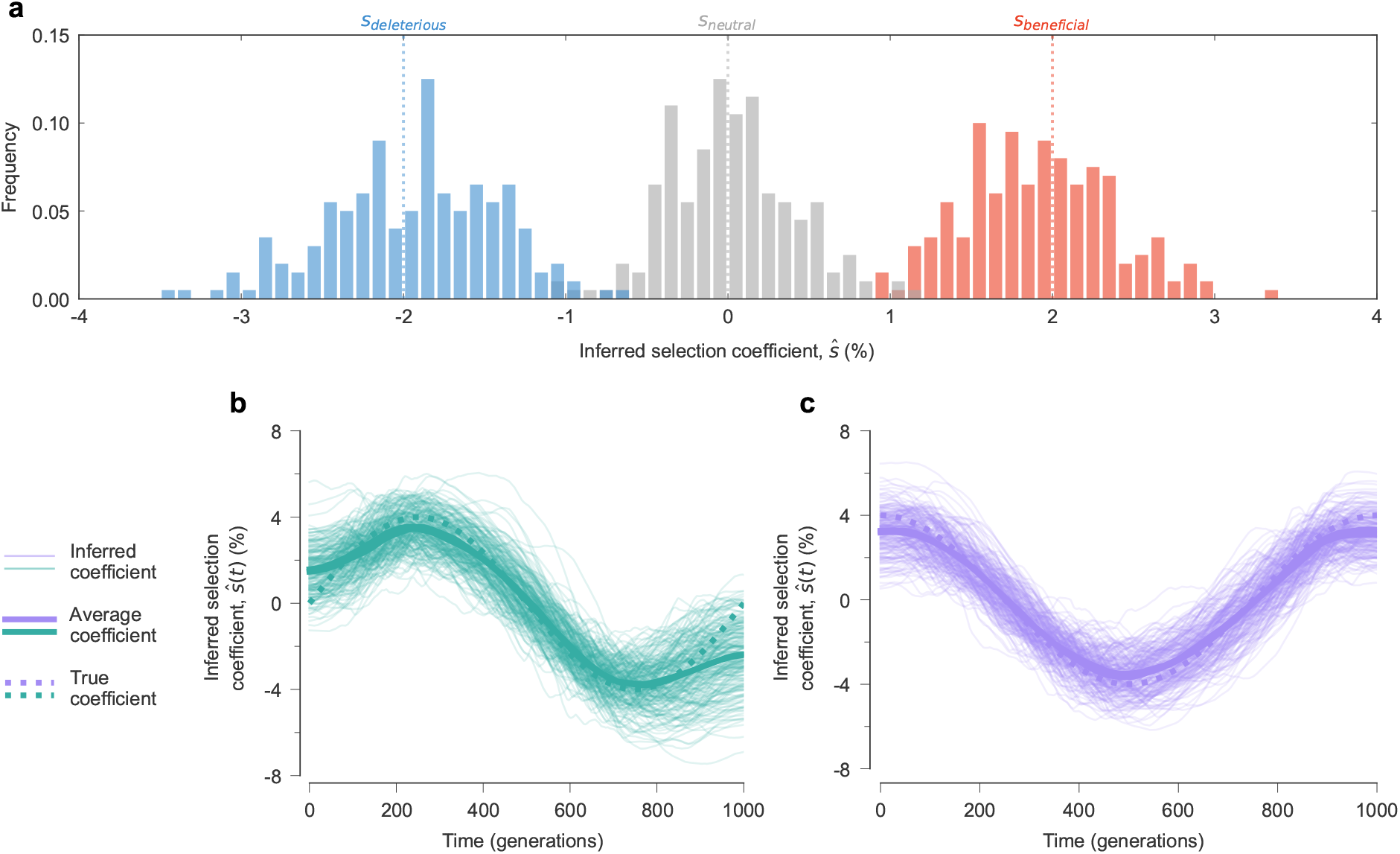
Robustness of inference across multiple replicate simulations. **a**, Distribution of constant selection coefficients inferred by our approach from 100 replicate simulations using the same parameters as in **Fig. 2. b, c**, The inferred trajectories of time-varying selection coefficients in individual simulations, and their average values, are close to the true ones.

We also systematically explored how different values of the temporal smoothness parameter *γ*′ affect our inference. We focused in particular near the boundaries of the observation window, where our inferences are less constrained by data (**Supplementary Fig. S2**; Supplementary Information). These tests helped us to identify appropriate choices for *γ*′that balance between overly rigid constraints (which might mask true fluctuations) and insufficient regularization (which could lead to spurious fluctuations, especially near the data boundaries).

### Fluctuating fitness shapes HIV-1 evolution

We applied our inference framework to analyze evolutionary data from HIV-1, focusing on viral adaptation within infected individuals. HIV-1 provides an ideal test case for our method because it evolves rapidly under strong, dynamical selection pressures from the host immune system. Although the immune response to HIV-1 is multifaceted, cytotoxic T lymphocytes (CTLs, also known as CD8^+^ “killer” T cells) play an especially important role in controlling viral replication ^33^. CTLs recognize and eliminate HIV-1-infected cells through a highly specific molecular mechanism (see **Fig. 4a**). Viral proteins inside infected cells are broken down into short peptide fragments, called epitopes, that are displayed on the cell surface by MHC molecules. CTLs can use their T cell receptors to detect these viral epitopes. This triggers the release of cytotoxic molecules that kill the infected cell, thereby limiting viral replication.

**Fig. 4.**
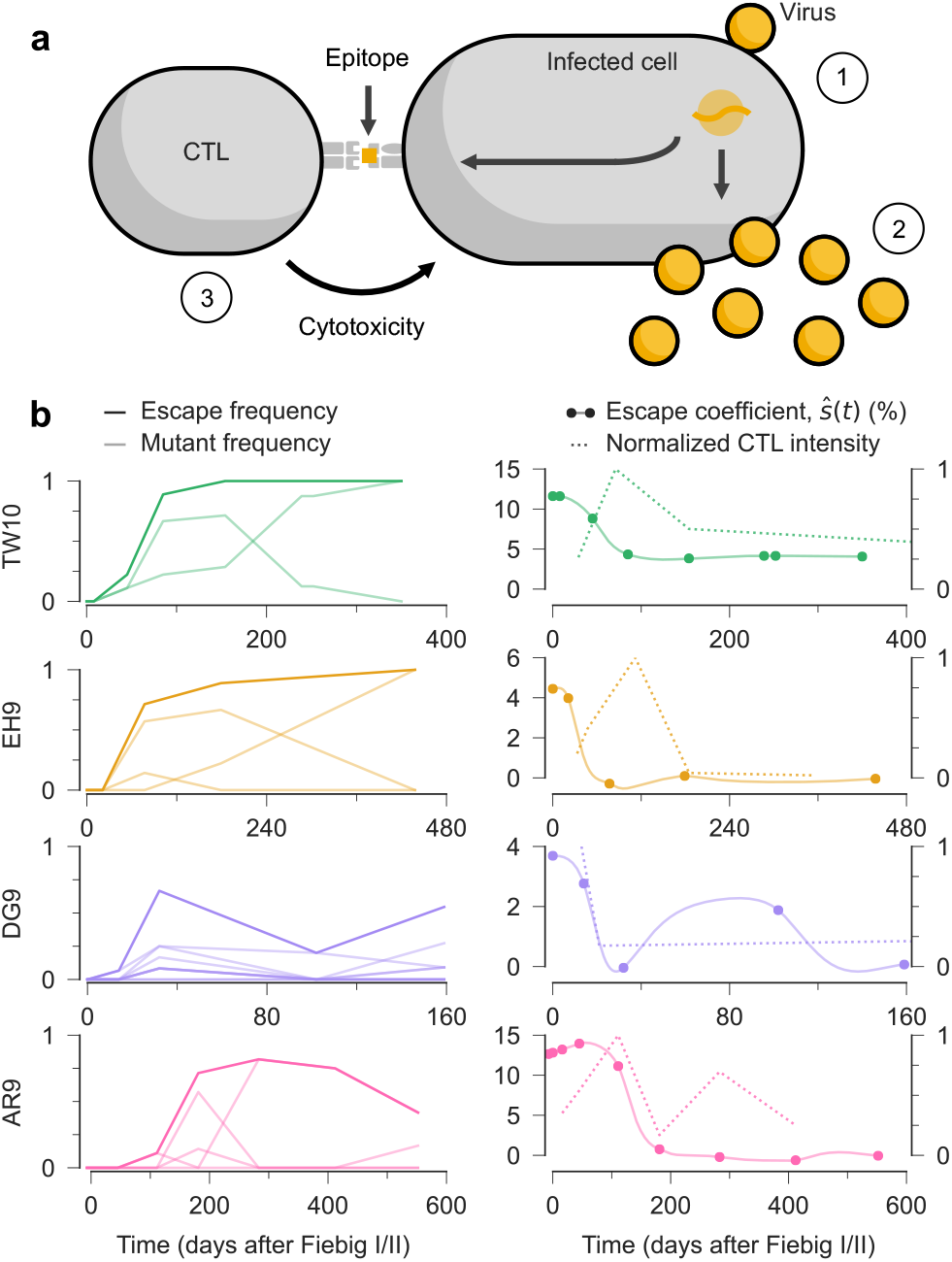
Estimates of time-varying selection for CTL escape. **a**, Schematic representation of HIV-1 infection and immune control. Viruses enter vulnerable cells (1) and replicate, producing new viral particles that can spread infection (2). However, CTLs can recognize viral epitopes presented on the infected cell surface, triggering the release of cytotoxic granules that kill the infected cell and cut short viral replication (3). **b**, Frequency of viruses with one or more escape mutations (escape frequency) and the frequency of individual escape mutations over time for four example CTL epitopes. Time is measured in days after Fiebig stages I/II, an early acute phase of HIV-1 infection roughly 1-2 weeks after transmission ^52^. The fitness benefit of escape that we infer (escape coefficient, *s* (*t*)) varies over time, peaking *before* the experimentally measured peak of the CTL response. For each epitope, CTL responses were normalized by dividing the measured spot forming units per million peripheral blood mononuclear cells (SFU/10^6^PBMCs) by its maximum value.

However, HIV-1 can escape immune control through mutations in CTL epitopes. Due to the virus’s high rate of mutation and replication ^34,35^, HIV-1 frequently acquires “escape mutations” that prevent CTL recognition while allowing the virus to continue replicating ^33,36–38^. This creates an evolutionary arms race, with HIV-1 evolution driven by the pressure to escape from dynamic immune responses.

Prior work has shown that the evolutionary pressure to escape CTLs is strong ^26,39–41^ but difficult to quantify precisely ^41–43^. A particularly puzzling observation is that viral escape from CTL responses that emerge later in infection tends to occur more slowly than escape from early responses ^44–46^. Several mechanisms have been proposed to explain this pattern, including competition between multiple escape variants (clonal interference) ^47^, greater fitness costs for escape mutations in later epitopes ^45^, and an increase in the breadth immune response ^48^. Here, we study how the fitness benefit of T cell escape shifts over time, compared with the dynamics of the immune response.

To capture the key biological features of immune escape, we extended our fitness model to differentiate between two types of selection pressures. First, we defined time-varying escape coefficients that quantify the fitness effect of having one or more mutations within a CTL epitope ^49^ (Supplementary Information). These coefficients capture the advantage of immune evasion. Second, we defined selection coefficients that quantify the effects of specific mutations (both within and outside CTL epitopes) on intrinsic viral replication, independent from immune evasion. We allowed escape coefficients to vary over time, reflecting changing immune pressures, while selection due to individual mutations was held constant. This follows from the biological intuition that immune selection likely fluctuates much more rapidly than other selective forces during infection. Tests in simulations confirmed our ability to distinguish these different fitness effects and track their changes over time (**Supplementary Figs. S3-S5**).

After validating our method in simulations, we applied it to longitudinal HIV-1 sequence data from 13 individuals ^50^. This data set offered several advantages. Donors were followed from early infection up to a few years before initiating antiretroviral drug therapy, capturing a critical period of immune-virus coevolution. Regular blood samples provided snapshots of the HIV-1 population over time as well as measurements of each individual’s immune response. In addition, the absence of antiretroviral drugs meant that immune pressure was the dominant force of selection guiding viral evolution. For our analysis, we incorporated HIV-1-specific rates of mutation ^51^ and recombination ^22^ to accurately model the evolution of the virus (Supplementary Information). To reduce confounding factors, we focused on 37 CTL epitopes where the fitness effect of immune escape could be statistically separated from the fitness effects of the underlying escape mutations and genetic background ^49^ (Supplementary Information). For 32 of these, the frequency and intensity of CTL responses against each epitope were also measured in enzyme-linked immunospot (ELISpot) assays ^50^. This allowed us to directly compare our estimates of selection with independent, experimentally measured immune activity.

### CTL intensity and the fitness effect of immune escape

Our analysis revealed a striking temporal pattern: for more than 90% of the CTL epitopes studied, the inferred fitness advantage of immune escape peaked at or before the maximum intensity of the corresponding CTL response (**Fig. 4b, Supplementary Figs. S6-S7**). This counterintuitive finding, where the selective advantage of CTL escape declines even as the immune response appears to become stronger, suggests an unappreciated complexity in HIV-1-immune dynamics.

This pattern could result from T cell exhaustion, a phenomenon in chronic viral infections where persistently stimulated T cells gradually lose their functional capabilities ^53–55^. T cell functions decline in a stepwise manner during exhaustion, and cytotoxicity is one of the first functions to be impaired ^53,54^. In contrast, secretion of the cytokine interferon gamma, measured by the ELISpot assays used in this study ^50^, is one of the CTL functions that is most resistant to exhaustion ^53,54^. Our findings therefore suggest that the actual cytotoxic pressure on the virus may have already begun declining by the time that cytokine production peaks. This could explain why the fitness benefit of escape decreases even while the strength of the immune response appears to increase.

### Kinetics of CTL escape

The evolutionary trajectories we observed varied significantly across different epitopes. For the majority of epitopes, we observed straightforward escape dynamics: once escape mutations appeared, they rapidly increased in frequency and remained dominant in the viral population for the duration of observations. However, a subset of epitopes exhibited more complex, non-monotonic patterns where escape variant frequencies fluctuated over time. Notable examples include the DG9 and AR9 epitopes (**Fig. 4b**), where escape frequencies increased initially before declining later in infection. The fitness benefit of escape that we infer reflects these dynamics, rising and falling along with escape mutants.

### Long-term decline in the fitness benefit of immune escape

Beyond the relationship with measured CTL responses, our analysis revealed a consistent long-term trend: the fitness advantage of immune escape declined substantially over the course of infection across nearly all epitopes (**Supplementary Fig. S8**). For many epitopes, selection for immune escape appears to be minimal a year or more after infection. This observation is aligned with previous observations of decreasing escape rates over time ^44–46^ but provides a more direct quantification of the underlying selection pressures. Because our model accounts for the intrinsic fitness effects of escape mutations, competition between escape variants, and correlations with mutations in the genetic background, these factors are unlikely to explain the declining benefit of immune escape.

However, our findings are consistent with the hypothesis of progressive immune dysfunction. In chronic infection, HIV-1-specific CTLs can become increasingly exhausted and lose their ability to effectively control viral replication. A reduction in CTL killing ability would also reduce selection for escape. This may explain why HIV-1 escape variants sometimes stall or even decline in frequency despite continued detection of the corresponding CTL response.

## Discussion

Evolution is guided by the shifting constraints of natural selection. Accurately quantifying these selection pressures represents both a major challenge and opportunity for evolutionary biology, with applications ranging from understanding adaptation in natural populations to guiding experimental evolution. Here, we developed a novel theoretical approach to infer a time-varying fitness seascape from genetic sequence data. Our approach is based on a path integral method from statistical physics, where “paths” represent histories of population evolution. Applying Bayesian inference methods, we found that the fitness effects of mutations that best explain an observed evolutionary history obey an Euler-Lagrange equation with the same mathematical form as the screened Poisson equation.

This connection to physics provides an intuitive interpretation. In our framework, changes in mutant frequencies that cannot be explained by mutation or recombination act as “charges” that reveal the underlying fitness seascape, analogous to the association between electric charges and the electric potential. The screening length in our equation determines how far in time these “charges” influence inferred selection, allowing us to distinguish between rapid fluctuations and more gradual shifts in fitness. Through extensive simulations, we confirmed that our method accurately recovers both constant and time-varying selection.

Applying our framework to longitudinal HIV-1 sequence data uncovered unexpected host-pathogen dynamics. We discovered that the fitness advantage of immune escape peaks at or before the maximum intensity of CTL responses, as measured by cytokine secretion assays. We also observed a consistent pattern of declining selection for immune escape over time across nearly all epitopes. This occurred despite the continued detection of HIV-1-specific CTL responses. Collectively, these observations are consistent with a model of progressive immune exhaustion in chronic HIV-1 infection. Chronically stimulated CTLs appear to lose their ability to control viral replication, weakening selection for immune escape. Our results highlight the dynamic interplay between viral evolution and immunity, showing that the fitness constraints guiding HIV-1 evolution can shift over time even within a single individual.

While our findings align with and extend prior work showing a reduced rate of immune escape late in infection ^44–46^, several limitations of the data should be considered when interpreting our results. Viral sequence data has inherent constraints. The limited number of sequences per time point (typically 5-20) and sometimes irregular sampling intervals introduce uncertainty in the true evolutionary trajectories. In turn, this increases the uncertainty in the fitness effects that we infer. CTL intensities were also sparsely sampled in time, making it difficult to track their dynamics precisely. Despite these challenges, the consistency of the relationship that we observe between selection for immune escape and CTL intensity supports the robustness of our results.

Future work could improve our inference framework. One limitation of our approach is that we treat the observed mutant frequencies (and their correlations) as exact, when in reality they are affected by finite sampling. Recent studies have developed clever methods to sample possible mutant frequency trajectories that are consistent with finitely-sampled data ^8,56^. Explicitly modeling sampling uncertainty could also improve the robustness of our results. This approach would be especially helpful for dealing with data that are sparsely sampled in time. Currently, we use linear interpolation between observed time points, which can occasionally introduce artefacts in inferred selection coefficients (see **Supplementary Fig. S9**). Though these effects appear relatively minor in our HIV-1 analysis, a more sophisticated approach that incorporates both sampling uncertainty and biologically plausible frequency trajectories between observations could yield even more robust inferences.

Overall, our work introduces a flexible theoretical framework for inferring fluctuating fitness seascapes from temporal genetic data. While demonstrated here using HIV-1 evolution, our method could be applied to study evolution in other biological contexts. Our framework can also readily be extended to incorporate more complex fitness functions (as shown in our analysis of HIV-1 immune escape) or features such as environmental covariates and spatial structure. This generality could enable new insights into diverse evolutionary phenomena in both experimental and natural populations. The ability to quantify how fitness landscapes shift over time represents an important step toward a more complete understanding of evolutionary processes in changing environments.

## ACKNOWLEDGEMENTS

The work of Y.G., B.L., and J.P.B. reported in this publication was supported by the National Institute of General Medical Sciences of the National Institutes of Health under Award Number R35GM138233.

## AUTHOR CONTRIBUTIONS

All authors designed research, performed research, and contributed new reagents/analytic tools. Y.G. analyzed data. Y.G. and J.P.B. wrote the paper.

## Supplementary Information

### Evolutionary model

In our study, we model the evolution of a population of *N* individuals subject to recombination, mutation, and natural selection, following the Wright-Fisher (WF) model. For simplicity, we first assume that each site can be in either a wild-type (WT) or mutant state. In population genetics, the terms “locus” and “allele” are often used to describe locations in the genome and the state of the genetic sequence there, respectively. For consistency with the evolutionary biology literature, we will use the same terminology. Individuals possess a genetic sequence length of 𝓁, resulting in a total of *M* = 2^𝓁^ possible genotypes. We further assume that there exist *λ* binary traits, which depend on the presence or absence of mutant alleles at specific sites and are subject to selection.

Let *n*_*a*_(*t*_*k*_) represent the number of individuals with genotype *a* at generation *t*_*k*_, and define *z*_*a*_(*t*_*k*_) = *n*_*a*_(*t*_*k*_)*/N* as the frequency of genotype *a* at generation *t*_*k*_. Thus, the state of the population at generation *t*_*k*_ can be defined by the vector *z*(*t*_*k*_) = (*z*_*a*_(*t*_*k*_), *z*_*b*_(*t*_*k*_), …, *z*_*M*_ (*t*_*k*_)).

In our fitness model, a positive selection coefficient *h*_*a*_ for genotype *a* indicates that this genotype has higher fitness compared to the wild type, whose fitness is set to 1. As in the constant inference case ^57^, we divide the fitness of each genotype into two parts: individual alleles (quantified by selection coefficients *s*_*i*_) and binary traits (quantified by trait coefficients *s*_*n*_). In the context of HIV-1 evolution, binary traits represent CTL escape: a virus either has mutations within an epitope, which we assume allows the virus to escape immunity, or it does not. We justify this approximation because the binding of T cell receptors to viral epitopes is highly specific ^57^. These traits could represent different biological features in other contexts. The fitness effects of different mutant alleles and binary traits are additive,

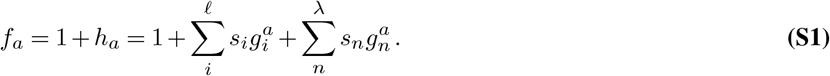

Here *g* is an indicator function to determine the presence of a mutation in locus *i* or trait *n*. We represent the haploid genetic sequence of each individual *a* as a binary string 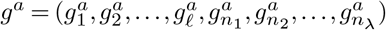 with 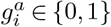. Here the index *i* labels each site in the genetic sequence, running from 1 to 𝓁; the index *n*_*i*_ labels each binary trait, running from *n*_1_ to *n*_*λ*_; and *a* labels the genotype (of which there are *M* = 2^𝓁^ possibilities). The effects of mutant alleles at different loci are cumulative, meaning that more beneficial alleles result in higher fitness. The fitness effects of different traits are also additive. In principle both the trait coefficients *s*_*n*_ and selection coefficients *s*_*i*_ can be functions of time, though we fix the selection coefficients *s*_*i*_ for our analysis of HIV-1 data.

The WF model also incorporates recombination and mutation. For simplicity, we assume that the recombination rate *r* and mutation rate *µ* are constant (though later we will relax this assumption). The mutation rate *µ* is the probability that the wild-type at one site converts to the mutant state, or vice versa. Given the exceedingly low mutation rate *µ*, we assume that each sequence undergoes at most one mutation per generation. Recombination refers to the process of genetic exchange between individuals, producing a “child” genome with shuffled contributions from each of the “parents.” We write the probability of a recombination breakpoint per site per reproduction cycle as *r*.

After accounting for the effects of selection, mutation, and recombination, the expected frequency of genotype *a* at generation *t*_*k*+1_, denoted as *p*_*a*_(*t*_*k*+1_), is

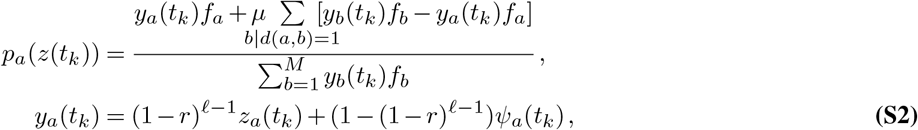

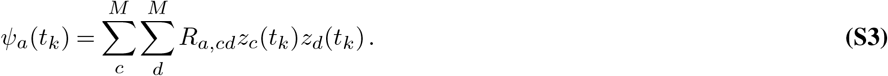

The notation *b*|*d*(*a, b*) = 1 indicates that genotypes *a* and *b* differ by just a single mutation. Under our assumption that each sequence undergoes at most one mutation each generation, genotypes *b* encompasses all genotypes that genotypes *a* can mutate to within one generation. The term *ψ*_*a*_(*t*_*k*_) represents the probability that the random recombination of any two individuals (*c* and *d*) within the population produces an offspring of genotype *a*, including cases where both parent and offspring share the same genotype *a*. Under WF dynamics, the probability of observing genotype frequencies ***z***(*t*_*k*+1_) at the next generation *t*_*k*+1_, given genotype frequencies of ***z***(*t*_*k*_)= (*z*_*a*_(*t*_*k*_), *z*_*b*_(*t*_*k*_), …, *z*_*M*_ (*t*_*k*_)) at generation *t*_*k*_, is multinomial:

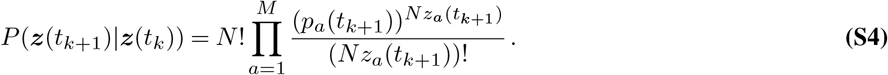

Consequently, given the selection and trait coefficients ***s***, the likelihood that the genotype frequency vector follows a specific evolutionary trajectory, (***z***(*t*_1_), ***z***(*t*_2_), …, ***z***(*t*_*K*_)), is

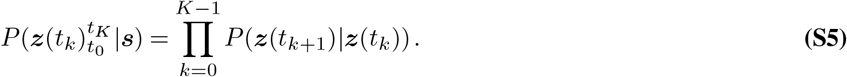

This complex likelihood function is conditioned upon the initial state ***z***(*t*_0_). Fortunately, (Eq. S4) can be simplified by using a diffusion approximation ^58^.

### Diffusion approximation

Since the populations are large and the changes in allele frequencies between generations are relatively small, we can transform the discrete, generation-by-generation model to a continuous-time model in the limit

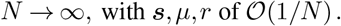

Given that the above parameters are of first-order infinitesimals (𝒪(1*/N*)), and dropping second-order and higher infinitesimals, *p*_*a*_(*t*) (S2) can be written as

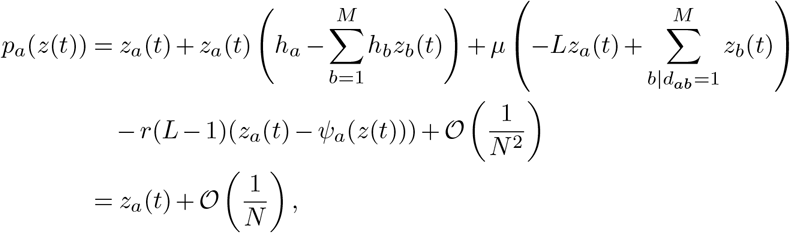

### Preliminary expansions for Δ*t* = 1

Since the Wright-Fisher model is a Markov process, we can use the following the conditional moment generating function

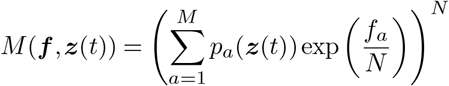

to obtain the following physical quantities:

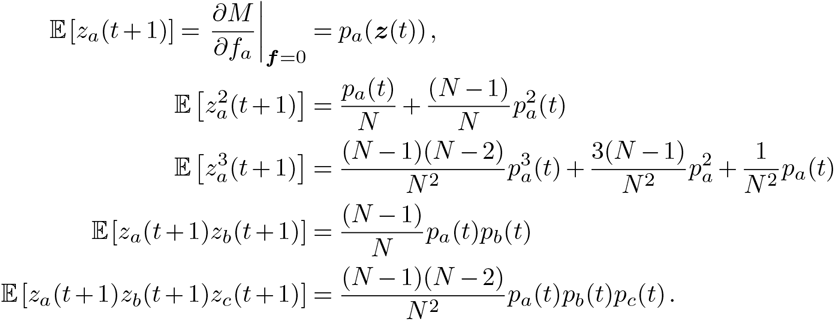

Based on these results, after neglecting second-order and higher infinitesimals, the first-order (drift), second-order (diffusion), and third-order terms for genotype frequency changes at Δ*t* = 1 are given by:

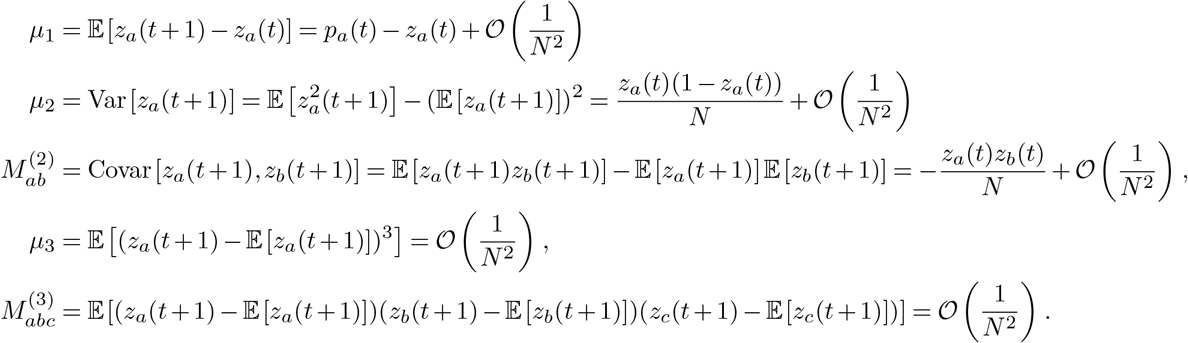

### Main derivations for arbitrary Δ*t*

To obtain the value at arbitrary Δ*t*, we incrementally advance *t* from 1 to Δ*t* in unit steps and apply mathematical induction to establish the result.

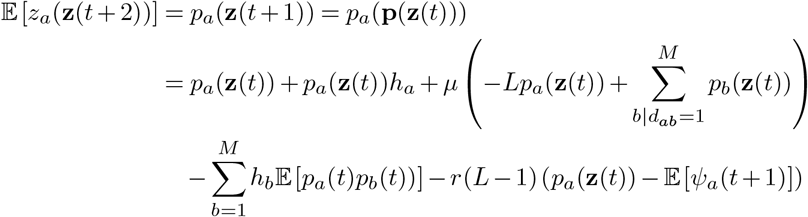

Here *ψ*_*a*_ is defined in S3. By applying the law of total expectation repeatedly, we can get the remaining expectations.

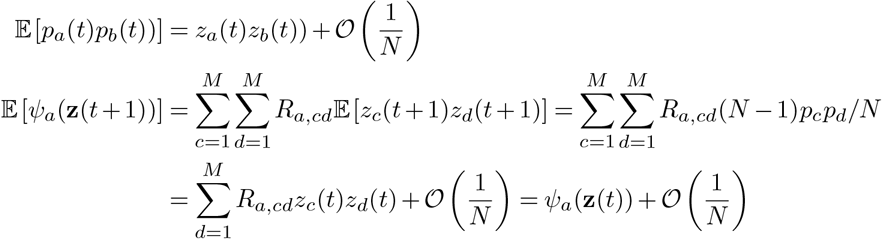

Thus,

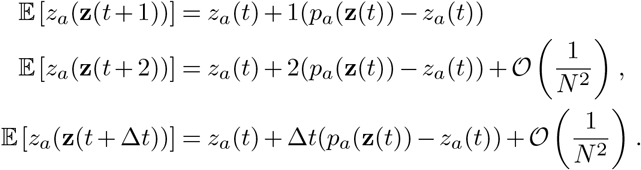

Because of the Markov property of the WF process, we can use the law of total expectation and the law of total variance to get

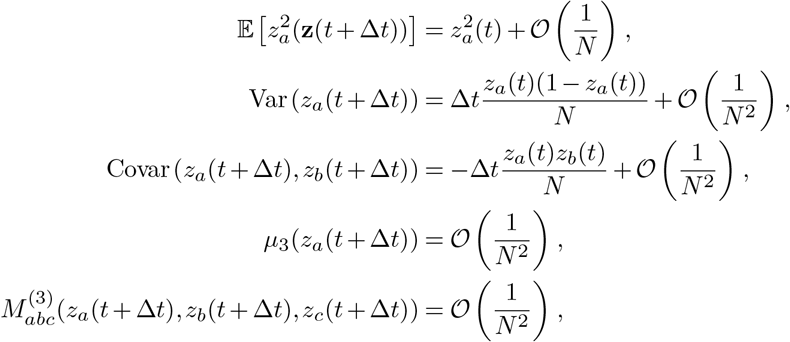

### Rescaling of time

In our framework, we can consider the scaling limit in which the population size becomes very large (*N*→ ∞), where frequencies transition from discrete values in the set 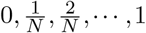 to a continuous values in the interval [0, 1]. We rescale time as 1*/N*, with the continuous difference *δt* = Δ*t/N*. After rescaling of time and frequency, we denote the genotype frequency as *ž*_*a*_(*t*). Considering Wright-Fisher model is a Markov process, and the increment *δž* of the process during a time interval [*t, t* + *δt*] satisfies the following conditions:

1. The expected value of Δ*ž*_*a*_ = *ž*_*a*_(*t* + *δt*) − *ž*_*a*_(*t*) is proportional to *δt*.

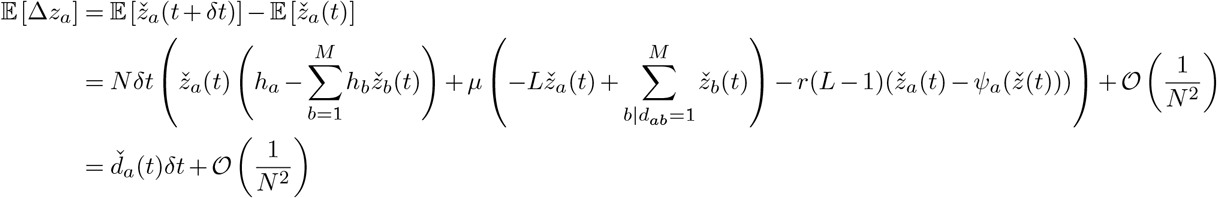

Here 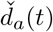 is known as the drift vector, describing the rate of expected changes in genotype frequencies at time *t* (see equation 4.100 of Risken ^59^). We will discuss its expansion in the next section.

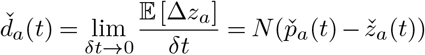
2. The variance of *δž*_*a*_ and covariance between *δž*_*a*_ and *δž*_*b*_ are also proportional to *δt*. Here we have

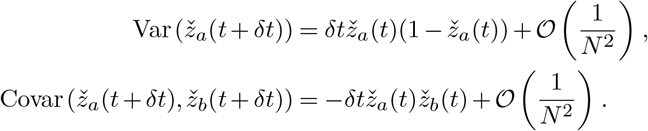

These terms are encapsulated in the diffusion matrix *Ď*_ab_(*t*), which describes the scaled covariance of the genotype frequency changes (see equation 4.100 of Risken ^59^).

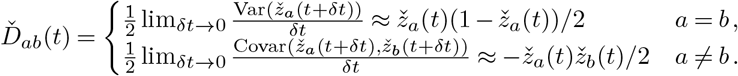
3. Higher order terms are subleading in 1*/N* and can be omitted. The random function *ž*_*a*_(*t*) is called a diffusion process ^58^. The diffusion process is described by the probability density function *ϕ*(***z***, *t*),

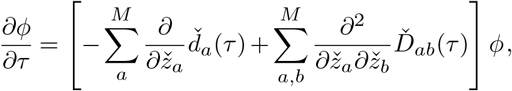

which is known as the Fokker-Planck equation.

### Path integral likelihood at genotype level

Under this diffusion approximation, using equation 4.109 of Risken ^59^, the transition probabilities (Eq. S4) become

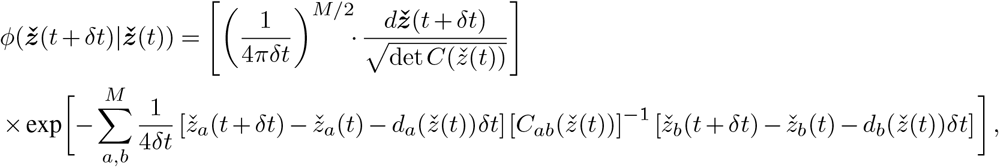

As noted before, here *d*_*a*_ are terms of the drift vector and *C*_*ab*_*/*2*N* are entries of the diffusion matrix. As seen from the definition, the covariance matrix *C* is a symmetric matrix with length *M*. At the genotype level, it can be written as:

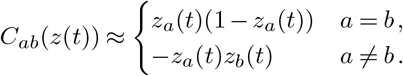

The drift vector is more complicated, and we will discuss its expansion later. This term is given by

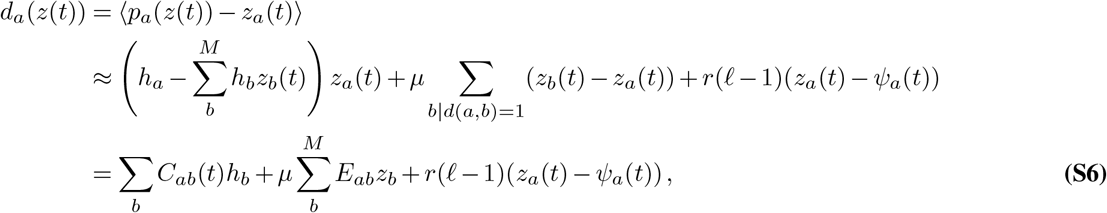

with the matrix *E*_*ab*_ reflecting the differences between genotype *a* and *b*,

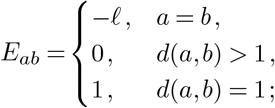

We can then express the likelihood of an evolutionary trajectory (Eq. S5) as a path integral, written in continuous time

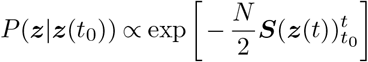

with the action 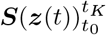 written as

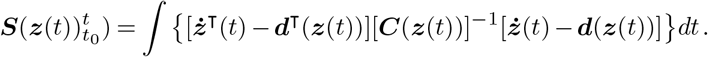

Here ***ż***(*t*) denotes the time derivative of the genotype frequency vector ***z***(*t*). In this expression, we have also written the frequency change, drift vector, and diffusion matrix in their vector/matrix forms, rather than explicitly writing genotype indices.

### Bayesian inference for time-varying selection

To control our estimates, we incorporate prior distributions for the selection and trait coefficients. Here we use a Gaussian distribution with zero mean for the selection coefficients ***s***. This approach helps curb the inference of strong fitness effects in the absence of strong statistical evidence:

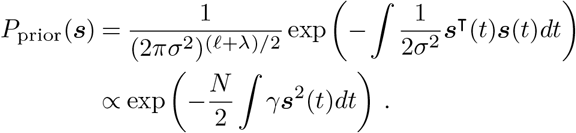

The maximum *a posteriori* estimate for the selection coefficients can then be found by maximizing the action *S*, similar to the case where selection coefficients are constant. However, this approach results in a problematic estimator that is extremely noisy with respect to time. This is because the estimates for ***s*** at different times are independent from one another, allowing estimates for successive times to vary dramatically. To address this issue, we add a Gaussian prior for the time derivative of the selection coefficients 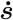, also with zero mean and 1*/*(*Nγ*′) variance. Combining these two prior distributions, we have:

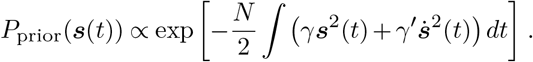

This prior implies that we expect the time-derivative of the selection coefficients to be small, that is, ***s*** changes slowly over time. By default, we set *γ* = 1, which slightly constrains magnitudes of inferred selection coefficients and helps to ensure that the matrix term is invertible. We set *γ*′ = 200 for time-varying selection coefficients. For constant selection coefficients, we can set a large *γ*′ (e.g., *γ*′ = 10^5^) to obtain a flat result.

With this prior distribution, the overall posterior distribution for the selection coefficients is then given by

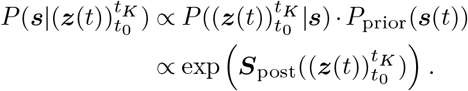

After applying Bayes’s theorem, the action 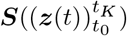 can be modified to 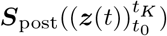, which can be written as

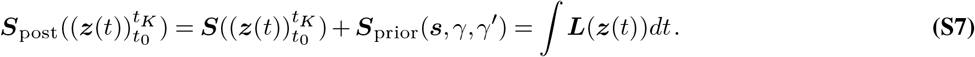

Here ***L*** is the Lagrangian function, with

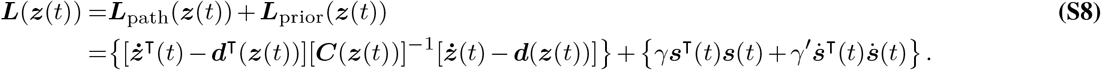

***d***(***z***) is the drift vector shown in Eq. S6. In the subsequent discussion, for brevity, we omit the time index *t*, as it is implicit for most physical quantities. All variables should be understood to be evaluated at *t* unless explicitly stated otherwise.

### Connection between allele level and genotype level

The maximum *a posteriori* selection coefficients can be found by maximizing the action (Eq. S7 and Eq. S8).

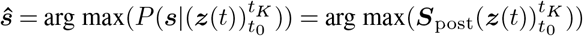

This equation contains information at both the genotype level (the genotype fitness gain, ***h*** and genotype frequency, ***z***) and the allele level (the selection coefficients ***s***). To unify the level of selection coefficients, we created a *M* × (𝓁 + *λ*) matrix ***G*** that bridges genotype and allele levels, with (*a, i*)th entries 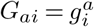 and (*a*, 𝓁 + *n*)th entries 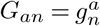. This matrix ***G*** allows us to express the relationship between genotype fitness gain ***h*** and selection coefficients ***s***. It also enables us to connect the genotype frequencies ***z*** with the allele frequencies ***x***.

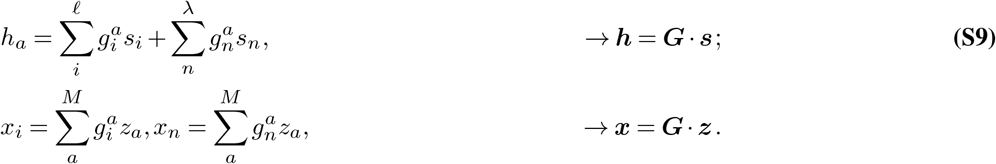

Here we denote the frequency of mutant alleles at locus *i* in the population as *x*_*i*_, and the frequency of individuals with one or more mutant alleles in binary trait *n* as *x*_*n*_. With the connection between ***h*** and ***s*** established through the matrix ***G***, we can rewrite the drift vector in Eq. S7 in terms of the selection coefficients ***s*** instead of the genotype fitness gain ***h***:

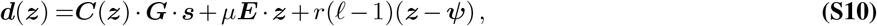

with 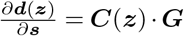. Here, we need to emphasize that the connection matrix *G* and the mutation index matrix *E* are both constant matrices and do not vary with time.

### Euler-Lagrange equation for the selection coefficients

Unlike the constant case, we applied the Euler-Lagrange equation 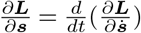 instead of simple differentiation to maximize the adjusted action (Eq. S7). This approach is necessary due to the time-varying nature of our selection coefficients. To obtain the results more conveniently, we take the transpose of both sides of the equation.

The right side of the equation equals

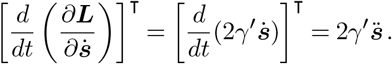

The left side of the equation is more complex. Given that the covariance matrix ***C***(***z***) is symmetrical, *C*^⊺^(***z***)[*C*(***z***)]^−1^ = *C*(***z***)[*C*(***z***)]^−1^ = *I*, where *I* is an identity matrix. LHS can be written as

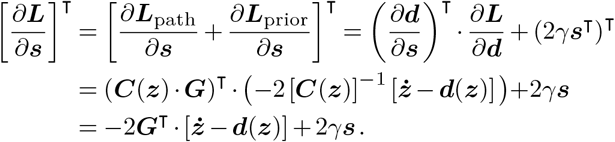

By equating these two sides, we obtain:

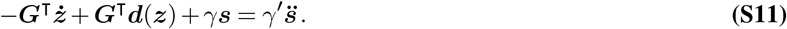

The results for the first term can be easily obtained. From equation S9, we can derive that 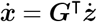. The second term is more complicated and requires further detailed analysis. We expand it as follows:

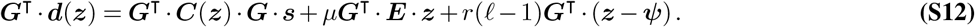

Our model consists of three main parts: the covariance matrix, mutation, and recombination. We’ll discuss each in turn. For covariance matrix part, we have

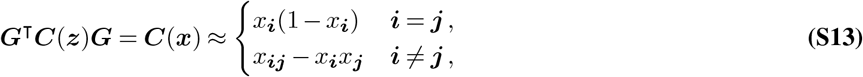

Here, ***i, j*** is a generic expression for locus *i* and binary trait *n*. We typically think of these binary traits as special virtual loci and use the generic index ***i, j***. It’s important to note that this index is distinct from the index of individual loci, which we denote as *i, j*.

Next, we examine the mutation term, which requires separate analysis for individual loci and binary traits. For the locus part:

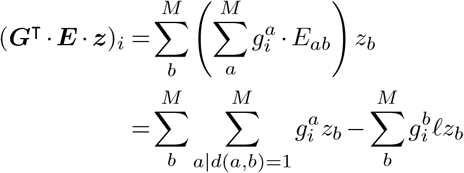

We can compute that 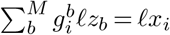. The first part requires a more detailed discussion of different scenarios. Since genotypes *a* and *b* differ by a single mutation, there are two scenarios:

1. 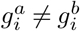: They differ at locus *i*, where each *b* only has one *a* satisfying the condition. In this scenario, 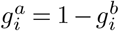.
2. 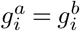: They differ at a locus other than *i*, where each *b* has (𝓁 − 1) corresponding *a*. This leads to:

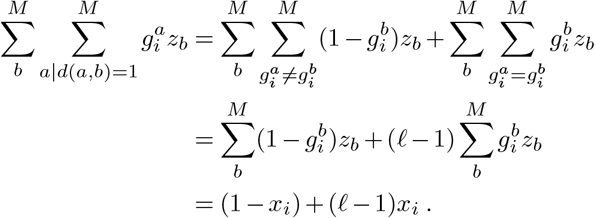

Thus, for locus part, the mutation term can be represented as

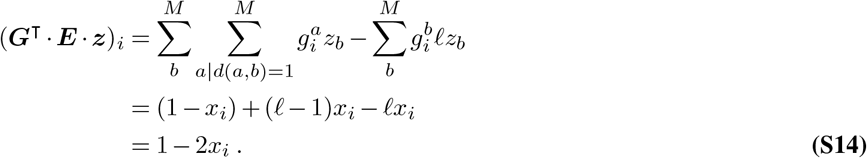

Using a similar method of case-by-case analysis, we can similarly get the conclude for binary trait part. For this part, given that genotype *a* and *b* differ by only one mutation, there are three scenarios:

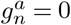: This then gives 0 directly regardless of the value of 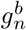.

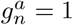 while 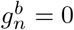: For the binary trait *n*, genotype *b* contains no mutation while genotype *a* has only one single mutation, where each genotype *b* corresponds to *l*_*n*_ = ∑_*i*∈*n*_ types of genotype *a*. This yields 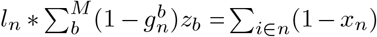.

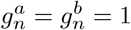: Both genotypes *a* and *b* have at least one mutation in binary trait *n*, which is a little complicated. We distinguish two additional cases based on the mutation number for *b* in binary trait *n*:

a. Single mutation in *b*: The *a* − *b* difference cannot be at *b*’s mutation locus. Each *b* corresponds to (𝓁 − 1) genotypes *a*. The probability for this case is 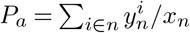, where 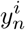 means the frequency of genotypes that contain only one mutation in trait group *n*, and the mutation is in locus *i*
b. Multiple mutations in *b*: The *a* − *b* mutational difference can be at any position. Each *b* corresponds to 𝓁 genotypes *a*. Thus, the results for this part are equal to: 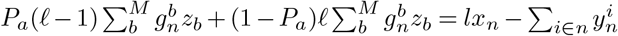.

Given the above, for binary trait part, the mutation term can be represented as

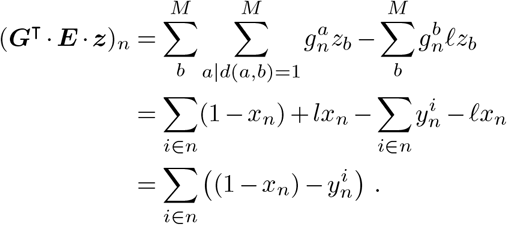

Interestingly, although the equations for individual loci and binary traits appear very different, we can discover that if we consider a single locus as a special binary trait, then the binary trait equation can be simplified to the locus equation. This serves as a verification of the correctness of our derivation for the trait part.

The recombination part is similar to the mutation part in its complexity. We begin with the locus part:

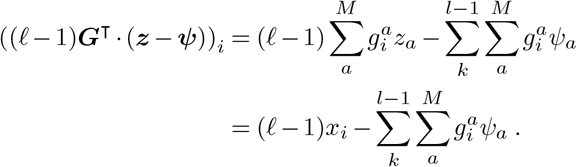

Here, let *k* index the 𝓁 − 1 possible breakpoints for a sequence of length 𝓁. To clarify the second term, we replace 𝓁 − 1 with its expanded form, the sum over all possible breakpoints.

Before discussing the different cases for the second term, we first define a new quantity 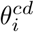

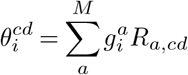

to represent the probability that the random recombination of any two individuals within the population results in an offspring with mutation at locus *i*. Here *R*_*a,cd*_ is described in equation S3. Thus,

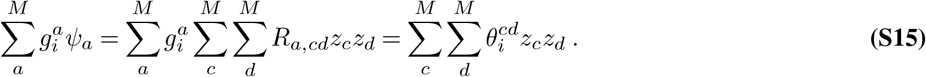

Based on the mutation status for genotype *c* and *d* at locus *i* (values of 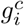 and 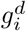), we have three scenarios:

1. No mutations 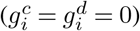: the offspring produced by recombination cannot have a mutation at *i*, so 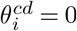.
2. Both mutated 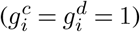: Offspring will have a mutation, 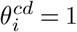.
3. Half mutated (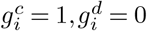 or 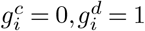): Half of offspring will have a mutation, 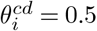.

Based on this, Eq. S15 can be represented as

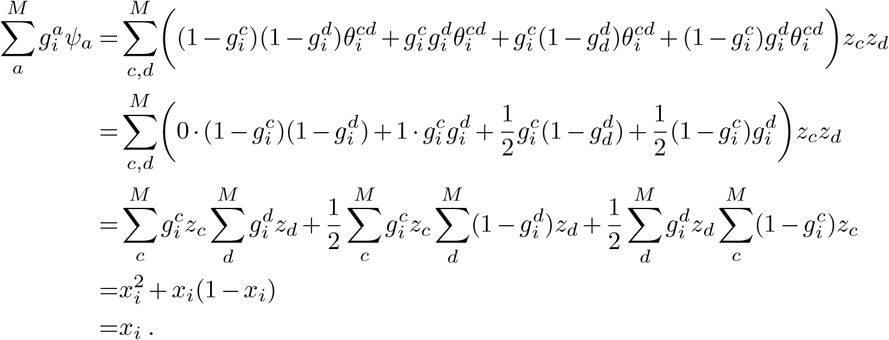

This leads to the conclusion that

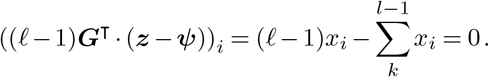

Now consider the binary trait part.

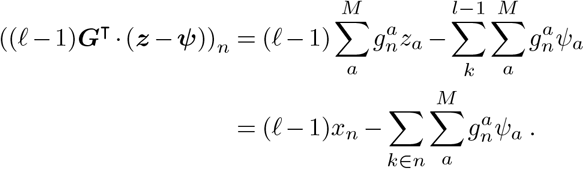

Similarly, we use 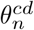 to represent the probability that recombination can result in an offspring with any mutation in binary trait *n*. Since a breakpoint can only alter the binary trait’s state if it falls within that trait, we restrict the breakpoint range to the binary-trait region rather than the entire sequence - using 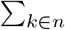 instead of 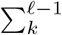. We again divide it into three scenarios:

1. No mutations 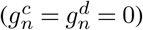: The offspring produced by recombination cannot have any mutation at *n*, so 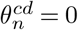.
2. Both mutated 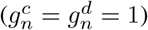: In most cases, the offspring will have mutations. However, a special case exists where half the offspring lack mutations. This occurs when, within a binary trait *n, c* has mutations only before break point *k*, and *d* only after *k*. Imagine two parental sequences, *c* and *d*, recombining at break point *k*. Focusing only on mutations within the binary-trait region, sequence *c* carries mutations exclusively upstream of k, while sequence *d* carries mutations exclusively downstream. Denoting mutated segments with c tilde (∼), we can write their sequences as: 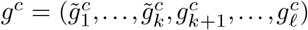 and 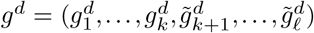. Then, the sequences for children *a* and *b* are 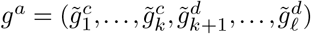 and 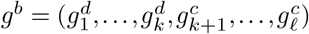. Here, we observe that although both *c* and *d* carry mutations (∼), child *a*, produced by recombining the non-mutated segments of *a*1 and *b*1, contains no mutations. The probability of the special case can be calculated as:

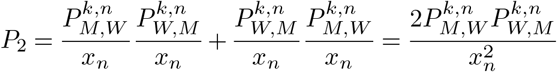

1. where 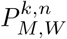 represents the frequency of sequences that have at least one mutant allele in the binary trait *n* before break point *k*, and all WT alleles in the same binary trait after *k*. With this probability, we can get 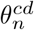 in this case:

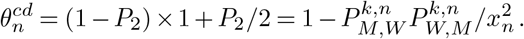

Half mutated (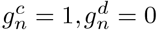 or 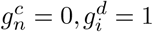): In most cases, half of the offspring will have mutations. However, a special case exists where all offspring have mutations. This occurs if the mutant sequence has mutations both before and after breakpoint *k*. (Parents: 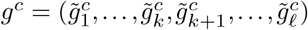 and 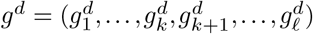. Children: 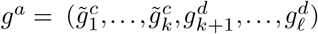 and 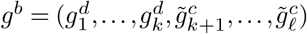.) The probability can be calculated similarly. Thus, in this scenario,

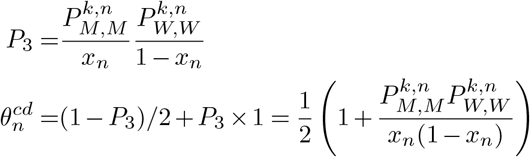

Thus we have

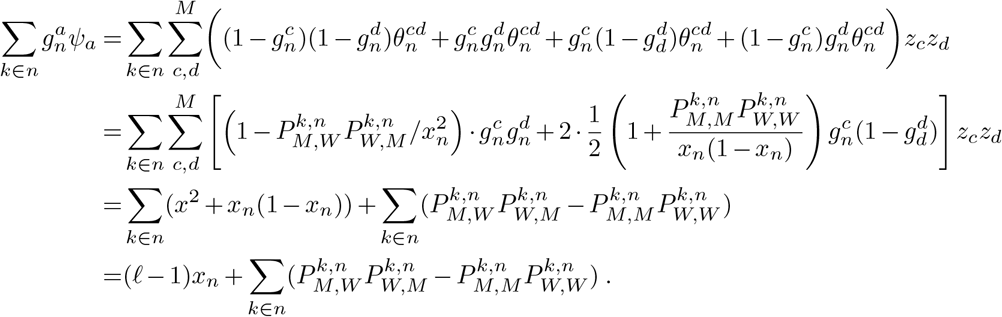

Considering all these scenarios, we can derive the final equation for the binary trait part:

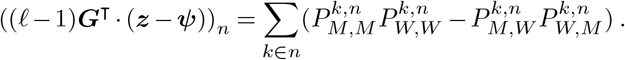

Let us now summarize the mutation and recombination terms here:

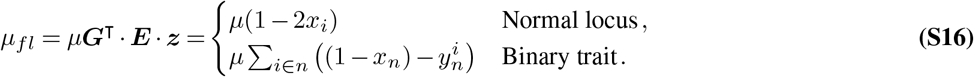

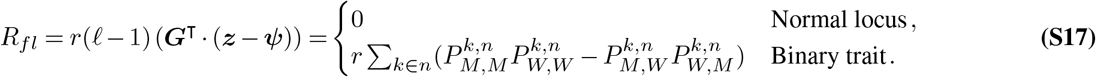

### Derivation of the differential equation

Building upon our previous derivations (Eqs. S11, S12, S13, S16, S17), we can express the equation in vector form:

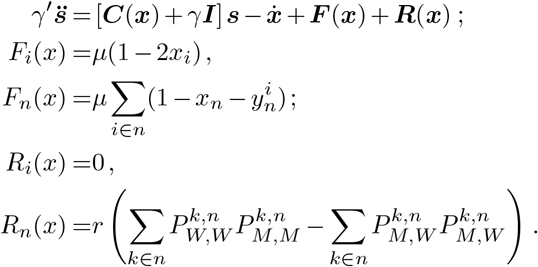

In this equation, ***I*** is the identity matrix, ***C***(***x***) denotes the allele/trait frequency covariance matrix, accounting for evolution speed correlations between mutations/traits, 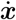 signifies the change in allele/trait frequencies at generation *t* and ***F*** (***x***) and ***R***(***x***) quantify the flux in allele/trait frequencies due to mutation and recombination separately.

Unlike the constant case, the best fit (maximum a posteriori) selection coefficients (Eq. S18) cannot be expressed analytically. This differential equation is second-order with time-varying coefficients (*C, F, R*) that lack algebraic expressions, precluding an analytical solution. To solve this differential equation, boundary conditions are necessary. However, there are no natural boundary conditions to impose on the selection coefficients ***s***, and their values at the endpoints are unknown. This scenario leads to free boundary conditions, resulting in the so-called transversality conditions 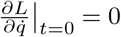 and 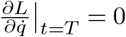, where *L* represents the Lagrangian. In our case, these conditions simplify to 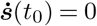 and 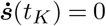.

### Extension to multiple alleles per locus and asymmetric mutation probabilities

To study real sequence data, we can extend the binary allele model presented in the previous sections to allow for multiple alleles per locus. In this section, we will derive results from an allele-level perspective while maintaining consistency with results derived from the genotype-level approach.

We use indices *α, β*,… to represent different alleles, with *q* denoting the total number of alleles (which could represent, for example, distinct nucleotides or amino acids). To accommodate both multiple-allele states for individual loci and binary states for traits, we adopt the following convention to maintain clarity in our notation: For individual loci, we employ the ordered pair (*iα*), where *i* denotes the locus and *α* specifies the allele at that locus. For binary traits, we retain the notation *n* for binary traits, which remains unchanged from the binary-allele case, as it naturally exists in only two states (WT and MT). The vector form for mutant allele frequencies at generation *t* can be written as a vector of length 𝓁𝓁 = 𝓁*q* + *λ*, encompassing all possible allele states at all loci and traits like this 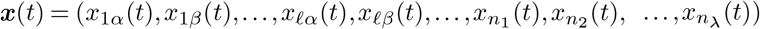.

The fitness model is expressed as:

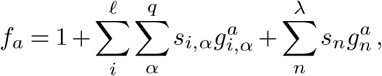

where *x*_*i,α*_ and *s*_*i,α*_ represent the frequency and selection coefficient for allele *α* at locus *i* respectively, and 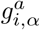 equals 1 if genotype *a* has allele *α* at locus *i*.

As before, we applied standard methods from statistical physics to convert the Fokker–Planck into a path integral that quantifies the probability density for “path” of mutant allele frequencies {***x***(*t*_1_), ***x***(*t*_2_), …, ***x***(*t*_*K*_)}. The probability for a path is proportional to the action 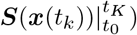, which is a function of allele frequency ***x*** instead of genotype frequency ***z***. Before Bayesian inference, the action can be expressed as:

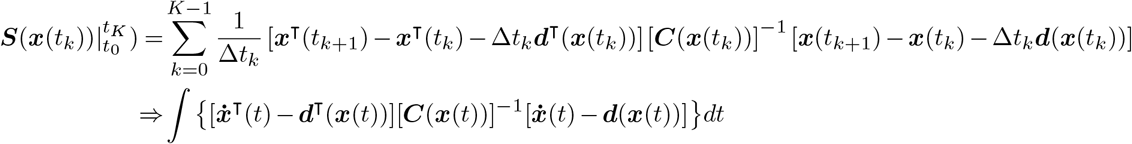

In this formulation, the drift vector ***d***(***x***(*t*)) and covariance matrix ***C***(***x***(*t*)) each comprise two distinct components. For simplicity in subsequent discussions, the time parameter *t* will be implicit unless explicitly required for clarity.

The covariance matrix ***C*(*x*)** is defined as an 𝓁𝓁 ×𝓁𝓁 matrix. After omitting terms of 𝒪(1*/N* ^2^), the covariance matrix can be expressed as:

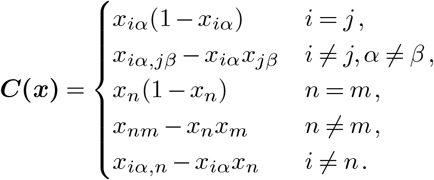

which is similar to ***C***(***z***). Here, *x*_*iα,jβ*_ represents the frequency of sequences with allele *α* and *β* occurring respectively at loci *i* and *j. x*_*iα,n*_ denotes the frequency of sequences with allele *α* at locus *i* and at least one mutation at binary trait *n*. The covariance between different alleles at the same locus is undefined, as each locus can only exist in one allele state at any given time.

Using previous equations (S6) and neglecting infinitesimals smaller than 𝒪(1*/N*), the drift vector ***d***_***i***_ is given by:

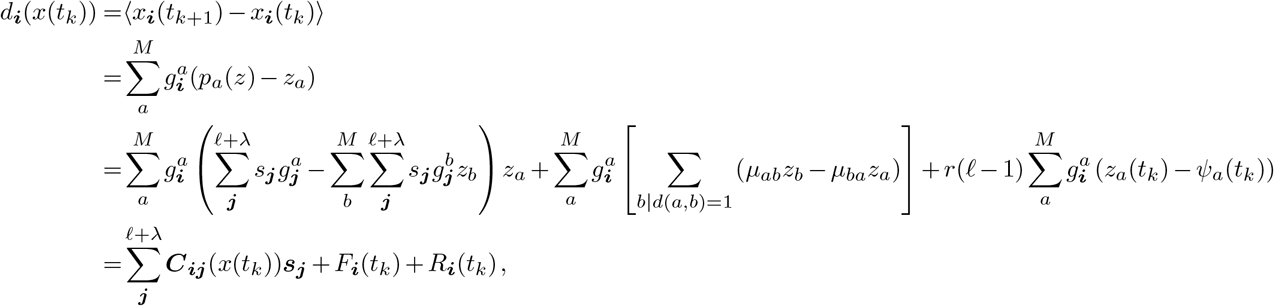

with

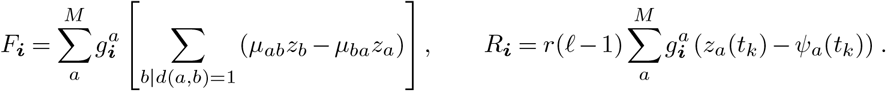

Here ***i*** is a generic locus, including pair (*i, α*) for individual loci and simple *n* for binary traits. This expression generalizes our previous formulations (Eq. S10) from the binary allele to the multiple allele case, incorporating the additional term *g*. This approach bridges allele-level and genotype-level perspectives. For the mutation term, we employ an asymmetric mutation matrix where the entry *µ*_*ab*_ represents the probability per locus per generation of mutation from genotype *a* to genotype *b*.

In the following discussion, we forgo a detailed derivation, instead focusing on the result and its physical interpretation. Since the binary trait part is not affected by the transition to the multiple allele case, and the recombination part for individual loci equals 0 (as recombination cannot alter individual allele frequencies), the recombination expression remains the same as before but requires careful consideration of synonymous mutations:

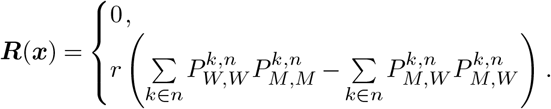

It’s crucial to note that synonymous mutant alleles do not contribute to binary traits and should be considered as wild-type alleles when calculating the recombination component for binary traits. The result here quantifies the net increase or decrease in mutant allele or binary trait frequency over time due to recombination.

The mutation component is more complex, requiring careful consideration of transitions between different alleles at the same locus. For an individual locus, the mutation component is expressed as:

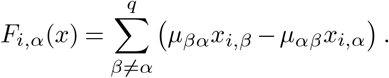

Here *µ*_*αβ*_ represents the probability per locus per generation of mutation from allele *α* to allele *β*. In this formulation, *µ*_*βα*_*x*_*i,β*_ quantifies the probability of allele *β* mutating to allele *α* at locus *i*. When summed over all possible alleles, this expression yields the net frequency change due to mutation. When *q* = 2 and assuming only one *µ*, this equation can be simplified to the binary case, given by Eq. S14.

The treatment of binary traits requires special consideration of synonymous mutations. Unlike individual loci, binary traits may have more than one “wild type.” We introduce *δ* to denote nonsynonymous mutant alleles, while *ϵ* denotes synonymous mutant alleles and wild type alleles. The mutation component for binary traits is expressed as:

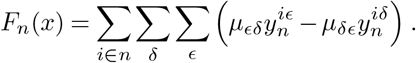

Here, 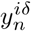 represents the frequency of sequences containing only one nonsynonymous mutation in trait group *n*, with mutation *δ* in locus *i*. This expression accounts for all loci within binary trait *n*, encompassing all possible nonsynonymous mutant alleles *δ* and all possible synonymous mutant alleles or wild type alleles *ϵ*. In the binary case, there is only one nonsynonymous mutation *δ*, and synonymous mutations *ϵ* can be seen as wild type. Thus, 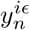 means the frequency for wild type in the binary trait *n*, which is (1−*x*_*n*_).

Our generalized formulation demonstrates consistency with simpler models through reduction to the binary-allele version under specific conditions. In the absence of synonymous mutant alleles within the binary trait, *y*^*iϵ*^ is the frequency of wild type for binary trait *n*, equal to 1−*x*_*n*_. When each trait only has a single *δ* and a single *ϵ*, our expression simplifies to match the binary-allele formulation. This reduction validates our approach, confirming its ability to encompass both simple and complex scenarios within a unified framework.

Thus, the final equation is

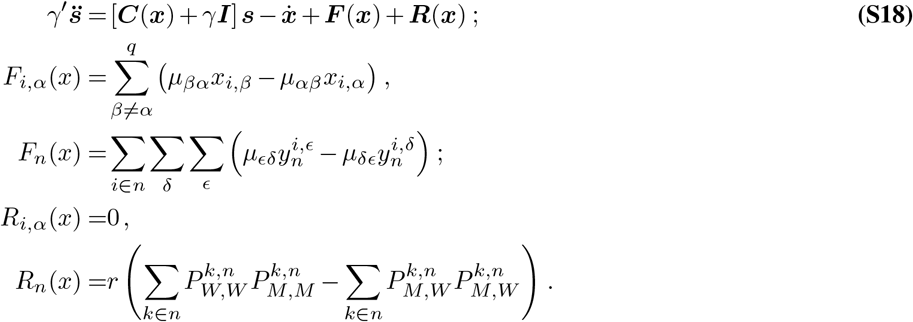

### Domain extension method

To apply the domain extension method to our equation, we first create a new extended domain 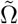. We extend the total generation by half in both forward and backward directions. The original time range in Ω spans from *t*_0_ to *t*_*K*_. In the extended domain 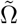, we add a segment of length 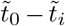 before *t*_0_, and a segment of length 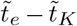 after *t*_*K*_, each equal to (*t*_*K*_ − *t*_0_)*/*2 (illustrated in **Supplementary Fig. S2**a). Then, we can set our extended transversality conditions:

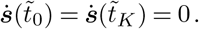

The source term in the extended domain 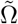 will be zeroed out. In addition, the covariance matrix will also be removed since there are no evolutionary forces outside the original region. Thus, the equation in 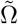 is only influenced by regularization force:

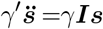

To ensure a smooth transition of the solution, we require continuity of the value of the solution and its derivative along the boundary ∂Ω:

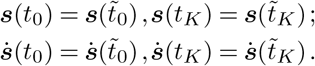

With all these equations and conditions, we can finally optimize the selection coefficients from *t*_0_ to *t*_*K*_.

### Testing performance in simulations

We conducted simulations of the time-varying Wright-Fisher model with discrete generations and binary (mutant/WT) states using Python. Our analysis encompassed two scenarios: a simple case without binary traits and a more complex case incorporating binary traits. For the simple case, we employed a simplified fitness model:

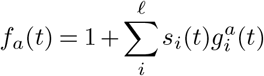

The complex case involved evolving populations of sequences according to our previously defined fitness equation (S1) over multiple generations.

For both cases, we started with an initial population of 4 random genotypes (each locus having a 20% probability of being mutant type). The value of *γ* is different from the value we used in constant inference. In constant inference, *γ* appears with 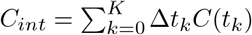, the sum of covariance matrix over all generations; while in the time-varying case, *γ* appears with *C*(*t*_*k*_), the covariance matrix at that generation. Terms in *C*_*int*_ and *C*(*t*_*k*_) can vary over several orders of magnitude, depending on the length of the trajectory. Thus, we need to use a smaller value regularization strength, such as *γ* = 10^−3^, to constrain the magnitude of the inferred time-varying selection coefficients. For the *γ*′ part, as in (**Supplementary Figs. S2** and **S5**), we extended the time range and used a time-varying value. The value we used for the boundary extended time is 4 times larger than the center time, which exponentially decreases to the middle value in a short time (10% of the total generations). For the center value, we set *γ*′ = 200 for time-dependent selection coefficients and *γ*′ = 10^6^ for time-independent selection coefficients. Parameter values are detailed in **Fig. 2** and **Supplementary Fig. S3** respectively.

### Influence of *γ*′ on inference

As discussed in the main text, the time coordinate in (2) practically behaves like a spatial coordinate in the context of inference. Similar to solving the electric field problem in Poisson’s equation, we have extended the range of generation *t* by adding half of the total generations to both the beginning and the end. For this outside area, all terms except for the two regularization terms due to the prior distribution will disappear, since we imagine there is no evolutionary force at those times. The value of *γ*′, which quantifies the expected time scale of environmental fluctuations, can influence the inference results. As described in the main text, 1*/*(2*γ*′) is the variance of 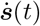, which means that a higher value of *γ*′ results in a smoother estimated result. At the boundary time points, we tend to use a large *γ*′ to avoid large fluctuations of selection or trait coefficient trajectories, which can change rapidly compared to times in the “bulk”, far from *t*_0_ and *t*_*K*_. Here we allocate 10% of of the total generations for *γ*′ to change its value, which is 100 generations in our simulations.

From repeated simulations (**Supplementary Figs. S2-S5**), we assessed how our method performs for different *γ*′ and could see a trade-off between average value and variance. A smaller *γ*′ means larger fluctuations and a larger variance of the inferred value (**Supplementary Fig. S2a**), while a larger *γ*′ can lead to a smoother inferred results (**Supplementary Fig. S2c**). When we have limited data, we tend to choose a larger *γ*′ at boundaries and that quickly reverts to normal values to prevent significant fluctuations at these points. However, in data sets with multiple replicates, a smaller *γ*′ typically yields an average value closer to the true value, particularly when the true value deviates significantly from 0. This comes at the cost of potentially extreme individual inference results.

We also observe that the root mean square error for the estimated selection coefficient is very large at the end of the trajectory. Even when we set a symmetric boundary condition for inference, the root mean square error is not symmetric. This is because we began with only 4 genotypes and ended with hundreds of genotypes, making the situation at the end of the trajectory much more complicated than the beginning.

### HIV-1 data

We obtained HIV-1 sequence data from 13 individuals of the CHAVI 001 and CAPRISA 002 studies in the United States, Malawi, and South Africa from the Los Alamos National Laboratory (LANL) HIV Sequence Database. For each individual, longitudinal HIV-1 half-genome sequences were collected from around the time of peak infection up to several months or years afterward. Donors did not receive antiretroviral drug treatment during this study.

Our data preparation process involved several steps to ensure data quality. We focused on heavily-sequenced half-genome regions, trimming full-length sequences accordingly. To minimize noise, we applied multiple selection criteria: We clipped full-length sequences down to the heavily-sequenced half-genome regions and applied several selection criteria to minimize the influence of noise in the data, including removing the sequences with large numbers of gaps, loci with high gap frequencies (*>*= 95% gaps), and time points with very small numbers of sequences (*<* 4) or large gaps (*>* 300) in time from the last sample. Following the removal of poorly sampled data points, ambiguous nucleotides were replaced by the most frequently observed nucleotides at the same site within the same individual. The final dataset consisted of 3′ and 5′ half-genome sequences of approximately 4,500 base pairs (bp) in length. Our analysis focused on polymorphic sites, defined as locations where more than one nucleotide (including gaps/deletions) was observed within an individual, typically around 100-900 bp in total. We designated the transmitted/founder (TF) sequence as the “wild-type” sequence for each individual. As noted above, synonymous mutant alleles within binary traits were considered “wild-type” for the purpose of computing the trait frequency.

The locations of CD8^+^ T cell epitopes in these sequences were determined both experimentally ^60^ and computationally ^61^. To isolate the fitness effects of escape from individual mutation effects, we focused on escape effects that could be independently inferred from other fitness contributions. This required that the escape trait be neither completely correlated nor anti-correlated with other variants. We accomplished this by reducing the integrated covariance matrix *C*_*int*_ = ∑_t_*C*(*t*_*k*_)Δ*t* to its reduced row-echelon form (RREF) and checking the linear dependencies of the rows for epitopes. The epitopes whose corresponding rows of the integrated covariance matrix are linearly independent are denoted as binary traits (ref. ^57,62^). These typically include epitopes containing three or more loci with non-synonymous mutations, though two loci may sometimes suffice. “Escape sites” are defined as polymorphic sites whose non-synonymous mutations were observed in the reading frame of an independent CD8^+^ T cell epitope. We anticipated that these non-synonymous mutations in escape sites would affect T cell recognition. Sites whose mutations can change the same epitope were considered part of a single “trait group.”

To infer selection, we used a mutation rate matrix ^63^ and a virus-load (VL) dependent recombination rate ^64^ as input. In our model, we dynamically determined the recombination rate based on the viral load at each time point (a simple linear model with *r* = 1.722 × 10^−10^ VL + 1.39 × 10^−5^), which was measured in past work ^60^. For later stages of infection where virus load was not measured, we assumed that VL values remained unchanged from the most recent measurement, consistent with the establishment of a viral set point in chronic infection.

We observed that the SR10 epitope in CH040 exhibited a relatively high escape frequency at the initial time point. To ensure biological plausibility of our results, we introduced a “negative time point” (−7 days) for CH040, roughly representing transmission, where all sequences were set to the TF sequence. This adjustment point can be observed in **Fig. 4b** and **Supplementary Fig. S7a**.

In our analysis, we employed *γ* to constrain the magnitudes of inferred coefficients, using different values for individual loci and binary traits: *γ* = 1*/* max(*t*) and *γ* = 10*/* max(*t*), respectively. Based on biological expectations, we also implemented asymmetric constraints. Specifically, we strongly expect that CTL escape should be beneficial for the virus, or at the very least, it should not be harmful to viral replication. We emphasize that this does *not* mean that individual escape mutations cannot be deleterious, or even that escape may be a net detriment to viral replication (that is, the benefit of escape may be outweighed by profoundly deleterious escape mutations). Rather, we claim that the property of being unrecognizable to CTLs on its own should not be harmful. Thus, the distribution of escape coefficients should not be symmetric around *s* = 0. Specifically, the variance in the *s <* 0 region should be smaller than that in the *s >* 0 region. Therefore, we applied a substantial penalty to negative escape coefficients by multiplying the corresponding *γ* values by 100. We also observed that once a mutation reaches fixation (*x* = 1), the *γ* parameter tends to drive the inferred selection coefficients toward zero due to a lack of evidence for persistent positive selection. To prevent the suppression of selection coefficients at later time points due to regularization effects alone, we implemented a smaller *γ* (1/10) after a mutation becomes fixed. For *γ*′, we used a constant value (*γ*′ = 10^6^) for time-independent selection coefficients and a time-dependent value (a similar pattern to **Supplementary Fig. S2c** in simulations, with a center value equal to 50, which is smaller than the value in simulations) for time-dependent coefficients, accounting for the sudden frequency changes characteristic of HIV-1 data.

## Data and code

Raw data and code used in our analysis are available in the GitHub repository located at https://github.com/bartonlab/paper-time-varying-selection. This repository also contains Jupyter notebooks that can be run to reproduce the results presented here.

---

**Supplementary Fig. S1.**
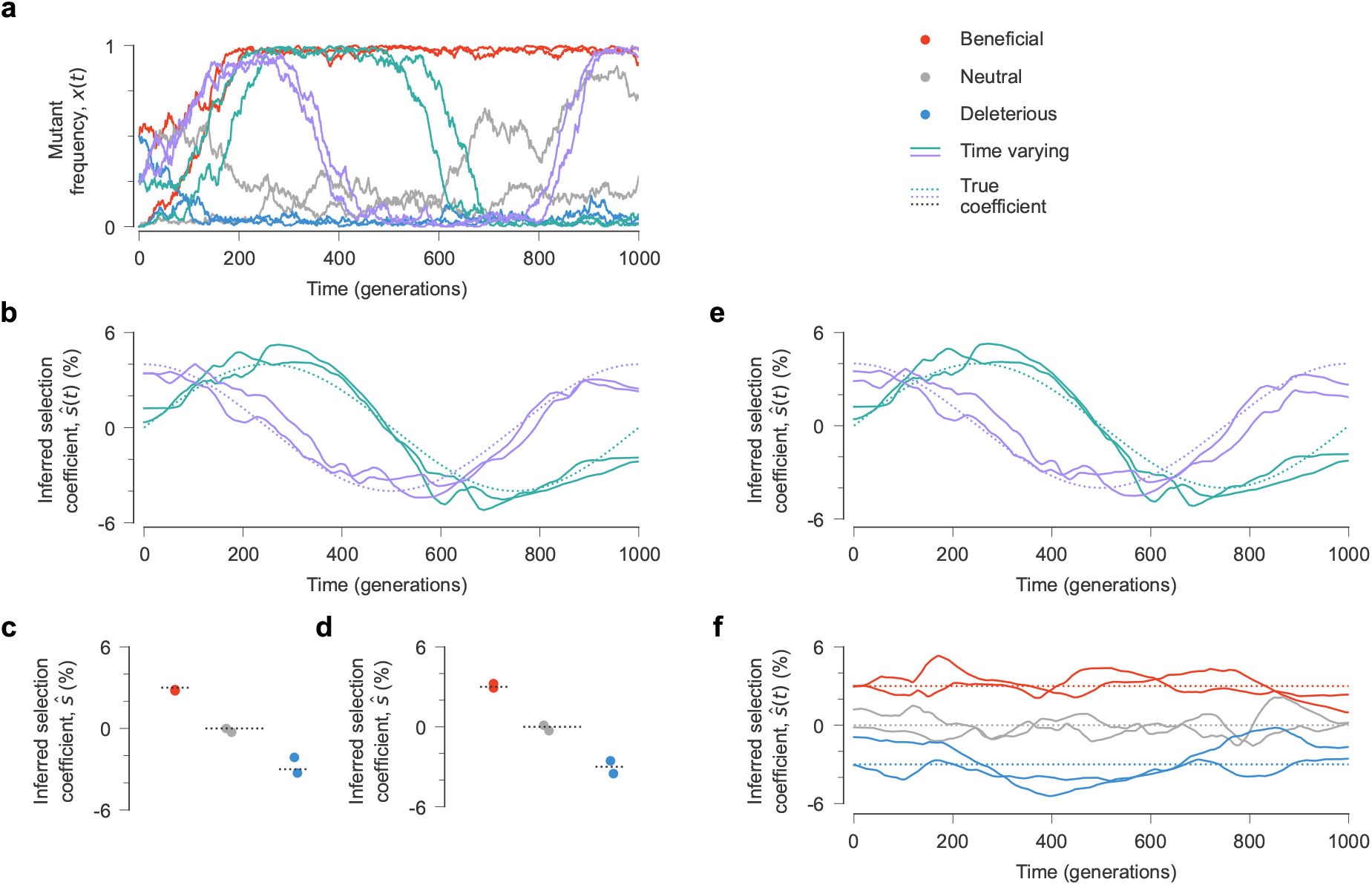
Robustness to model misspecification. **a**, Simulated mutant allele frequency trajectories. Time-varying selection coefficients (**b**) and constant ones (**c**) inferred with prior knowledge of the constant sites. To test the robustness of our approach to model misspecification, we also tested a scenario where all selection coefficients were assumed to be time-varying, including the constant ones. In that case we obtained time-varying coefficients (**e**) that were very similar to the previous ones. Inferred selection for the constant coefficients (**f**) fluctuated around the true, underlying constant values. To help visualize the long-term average, we plotted the mean of these inferred selection coefficients over time (**d**) versus their true, constant values. Simulation parameters: 𝓁 = 10 loci with two alleles at each locus, two beneficial mutants with *s* = 0.03, two neutral mutants with *s* = 0 and two deleterious mutants with *s* = −0.03. Additionally, four mutations have fitness effects that vary over time, with two following a sinusoidal pattern (green) and the others a cosine pattern (purple). The mutation probability per site per generation is *µ* = 1*×*10−3, the recombination probability per site per generation is *r* = 1*×*10−3, and the population size is *N* = 10^3^. The initial population was randomly generated and evolved over *T* = 1000 generations.

**Supplementary Fig. S2.**
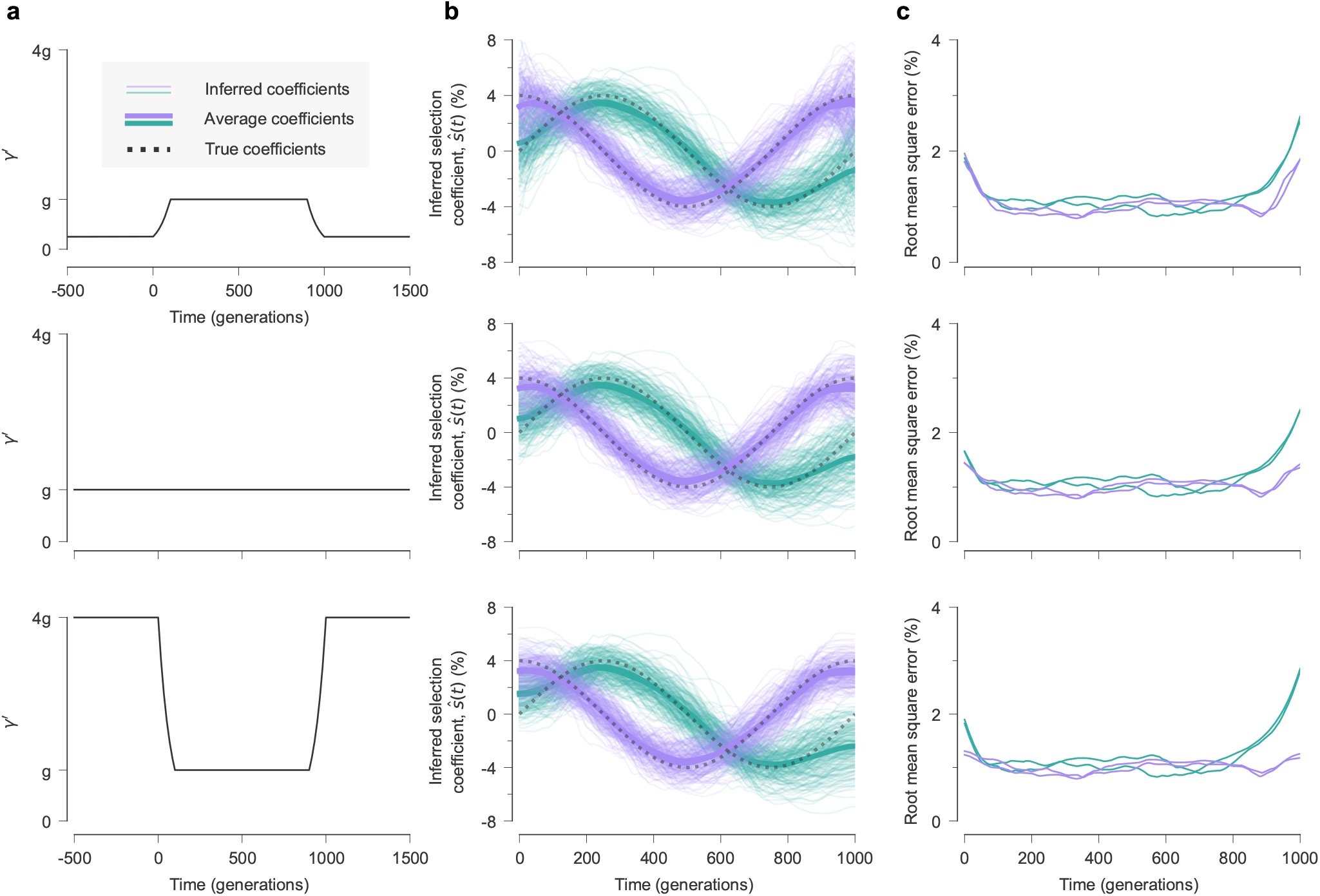
Effects of different *γ′* profiles on inference near the boundary. In real data and simulations, the data that we use for inference is bounded between some initial time *t*_0_ and final time *t*_K_ (here zero and 1000, respectively). Selection coefficients near *t*_0_ and *t*_K_ are less well-constrained than those in the “bulk” far from the time boundaries because we receive no information from times before *t*_0_ or after *t*_K_. Here, we examined how changes in *γ′* in the bulk and beyond the boundaries affect our inference. Column **a** shows the change of *γ′* across time. Time before 0 and after 1000 are the extended time. Column **b** displays the inferred time-varying selection coefficients and their averages across 100 independent simulations with identical underlying fitness parameters. Column **c** shows the root mean square error of inferred values across 100 simulations as a function of time. The inferred coefficients closely match the true values in the central time region across all cases. However, when *γ′* takes small values near the extended boundaries, the inferred coefficients exhibit significant fluctuations in these regions, with a higher RMSE. Increasing *γ′* near the boundaries suppresses fluctuations, which can sometimes decrease inference errors. Simulation parameters: 𝓁 = 10 loci with two alleles at each locus (mutant and wild-type, WT), two beneficial mutants with *s* = 0.02 (red), two neutral mutants with *s* = 0 (grey), and two deleterious mutants with *s* = −0.02 (blue). We consider four mutations with time-varying fitness effects, with two following a sinusoidal pattern (green) and the others a cosine pattern (purple). The mutation probability per site per generation is *µ* = 1*×*10^−3^, the recombination probability per site per generation is *r* = 1*×*10^−3^, and the population size is *N* = 10^3^. The initial population was randomly generated and evolved over *T* = 1000 generations, with each locus having a 20% probability of being mutant type.

**Supplementary Fig. S3.**
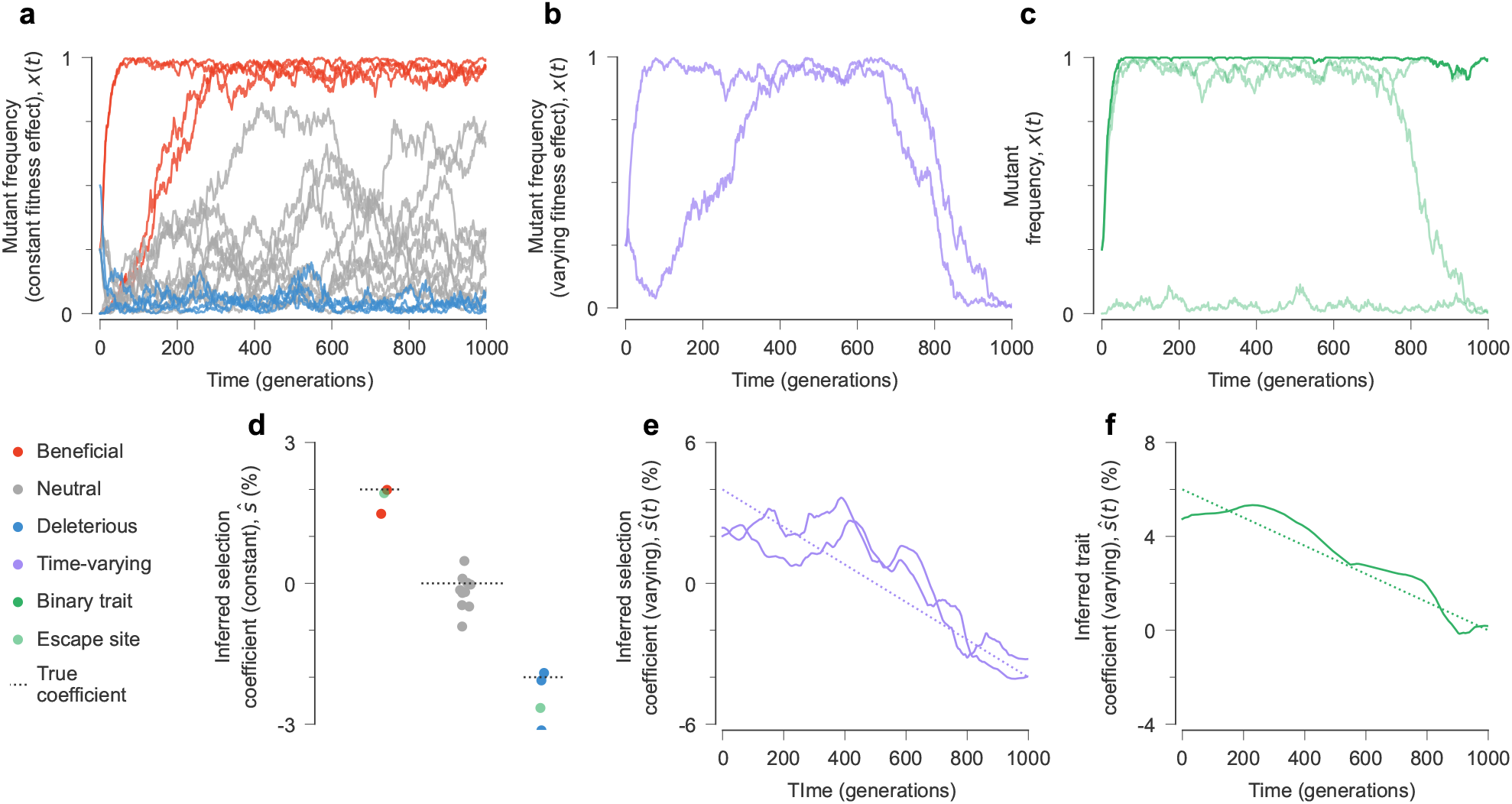
Inferring selection on binary traits with time-varying fitness effects from temporal genetic data. **a**, Frequency trajectories for alleles with constant fitness effects. **b**, Frequency trajectories for alleles with time-varying fitness effects. **c**, Trait frequencies and their contributing individual allele frequencies in the same simulation. The inferred constant selection coefficients (**d**), time-varying selection coefficients(**e**), and time-varying trait coefficients (**f**) are close to their true values. Simulation parameters: 𝓁 = 20 loci with two alleles at each locus (mutant and wild-type, WT) including several mutations with constant fitness effects (beneficial mutants with *s* = 0.02 (red), neutral mutants with *s* = 0 (gray), deleterious mutants with *s* = −0.02 (blue) and mutants contributing to the binary trait (green)) and two mutations with time-varying fitness effects. Green mutations have two distinct effects on fitness: one through their individual selection coefficients (**d**) and one through their contribution to the time-varying trait effect (**e**), analogous to immune escape in the context of HIV-1. The mutation probability per site per generation is *µ* = 1*×*10^−3^, the recombination probability per site per generation is *r* = 1*×*10^−3^, and the population size is *N* = 10^3^. The initial population was randomly generated and evolved over *T* = 1000 generations, with each locus having a 20% probability of being mutant type.

**Supplementary Fig. S4.**
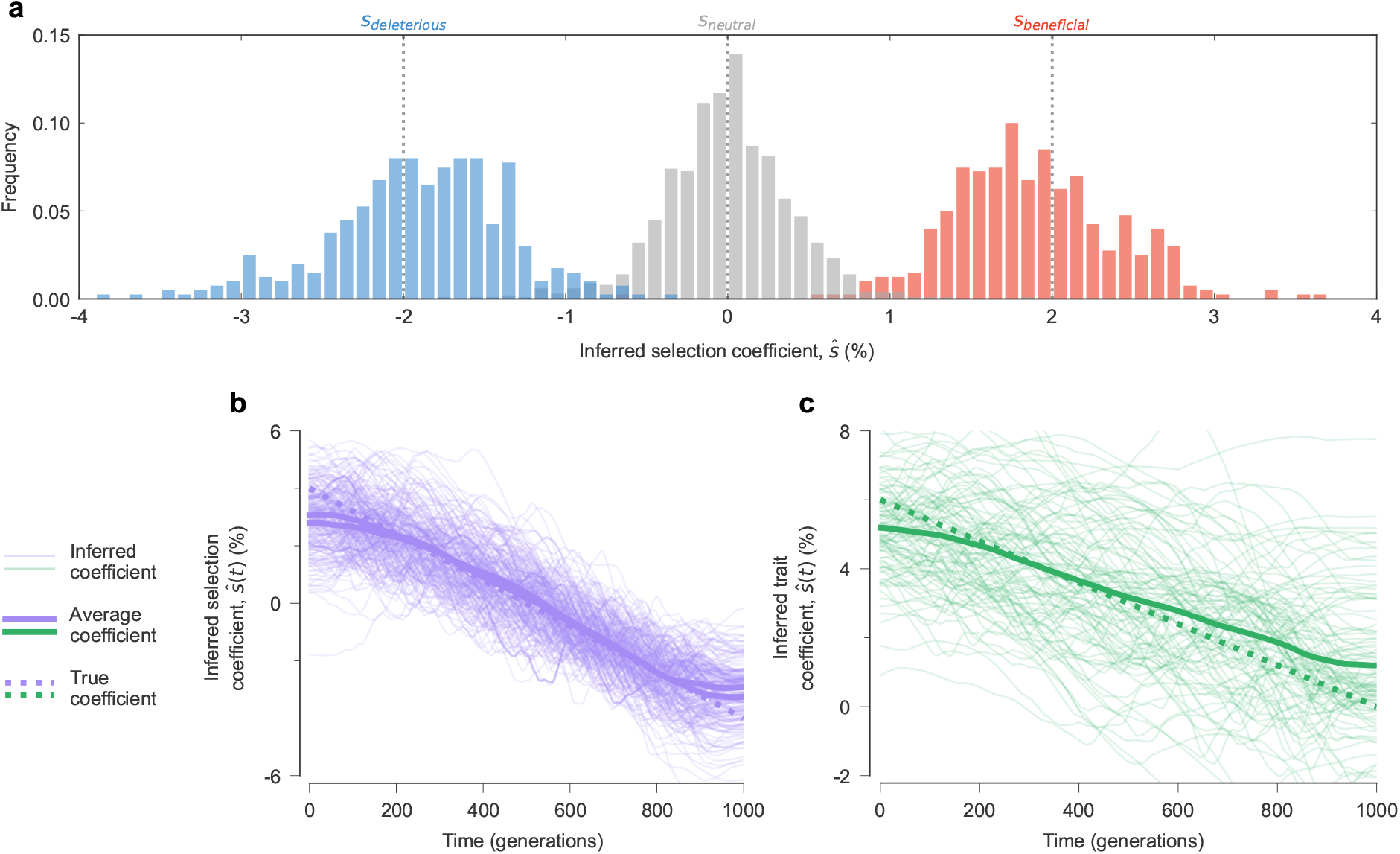
Consistency of inference across multiple replicate simulations including time-varying fitness effects of individual mutations and binary traits. Distribution of **a** inferred constant selection coefficient estimated across 100 replicate simulations, using the same initial parameters. **b** and **c** show the inferred time-varying selection coefficients for individual mutant alleles and the binary trait, respectively. Simulation parameters: 𝓁 = 20 loci with two alleles at each locus (mutant and wild-type, WT) including several constant mutations (beneficial mutants with *s* = 0.02 (red), neutral mutants with *s* = 0 (gray), deleterious mutants with *s* = −0.02 (blue)) and two time-varying mutations (purple)). Sites contributing to binary traits (green) in these 100 simulations were randomly selected from sites with constant selection coefficients. The mutation probability per site per generation is *µ* = 1*×*10^−3^, the recombination probability per site per generation is *r* = 1*×*10^−3^, and the population size is *N* = 10^3^. The initial population was randomly generated and evolved over *T* = 1000 generations, with each locus having a 20% probability of being mutant type.

**Supplementary Fig. S5.**
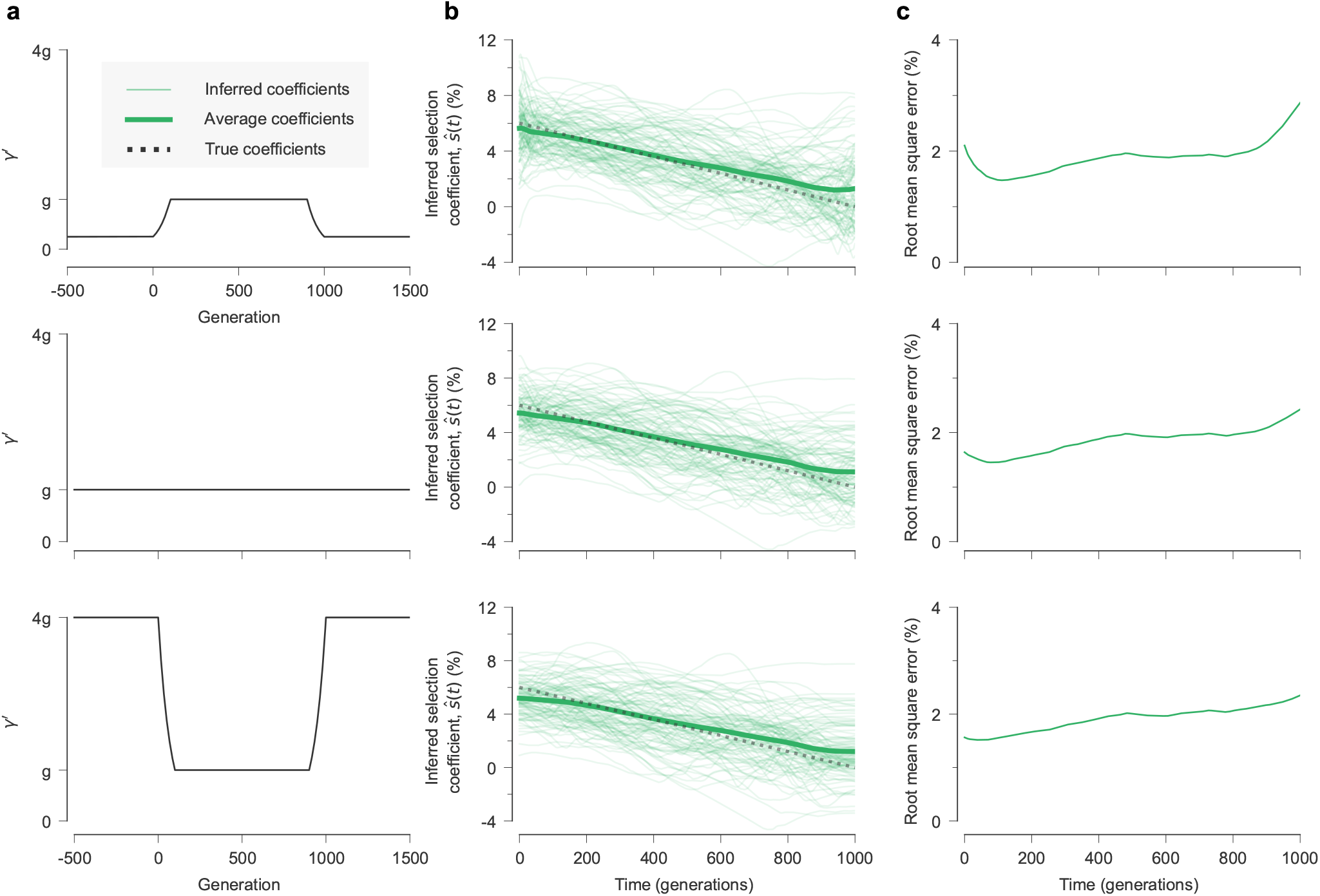
Effects of different *γ′* profiles for inference of selection on binary traits near the time boundaries. This figure is analogous to **Supplementary Fig. S2**, but here we explore the effect on trait inference. **a**, Profiles of *γ′* over time. Times before 0 and after 1000 are in the extended domain, beyond the bounds of the simulation. **b**, Inferred time-varying trait coefficients and their average values across 100 independent simulations using identical initial parameters. Here we used the same simulation parameters as in **Supplementary Fig. S3. c**, Root mean square error of inferred values across 100 simulations as a function of time. The inferred coefficients closely match the true values in the central time region across all cases. However, when *γ′* uses small values near the extended boundaries, the inferred coefficients exhibit significant fluctuations in these regions, with a higher RMSE. Increasing *γ′* around the boundaries suppresses fluctuations due to the lack of constraints in the extended domain. Simulation parameters: 𝓁 = 20 loci with two alleles at each locus (mutant and wild-type, WT) including several constant mutations (beneficial mutants with *s* = 0.02, neutral mutants with *s* = 0, deleterious mutants with *s* = −0.02), three random mutants that contribute to the binary trait, and two mutations with time-varying fitness effects. The mutation probability per site per generation is *µ* = 1*×*10^−3^, the recombination probability per site per generation is *r* = 1*×*10^−3^, and the population size is *N* = 10^3^. The initial population was randomly generated, with each locus having a 20% probability of being mutant type, and evolved over *T* = 1000 generations.

**Supplementary Fig. S6.**
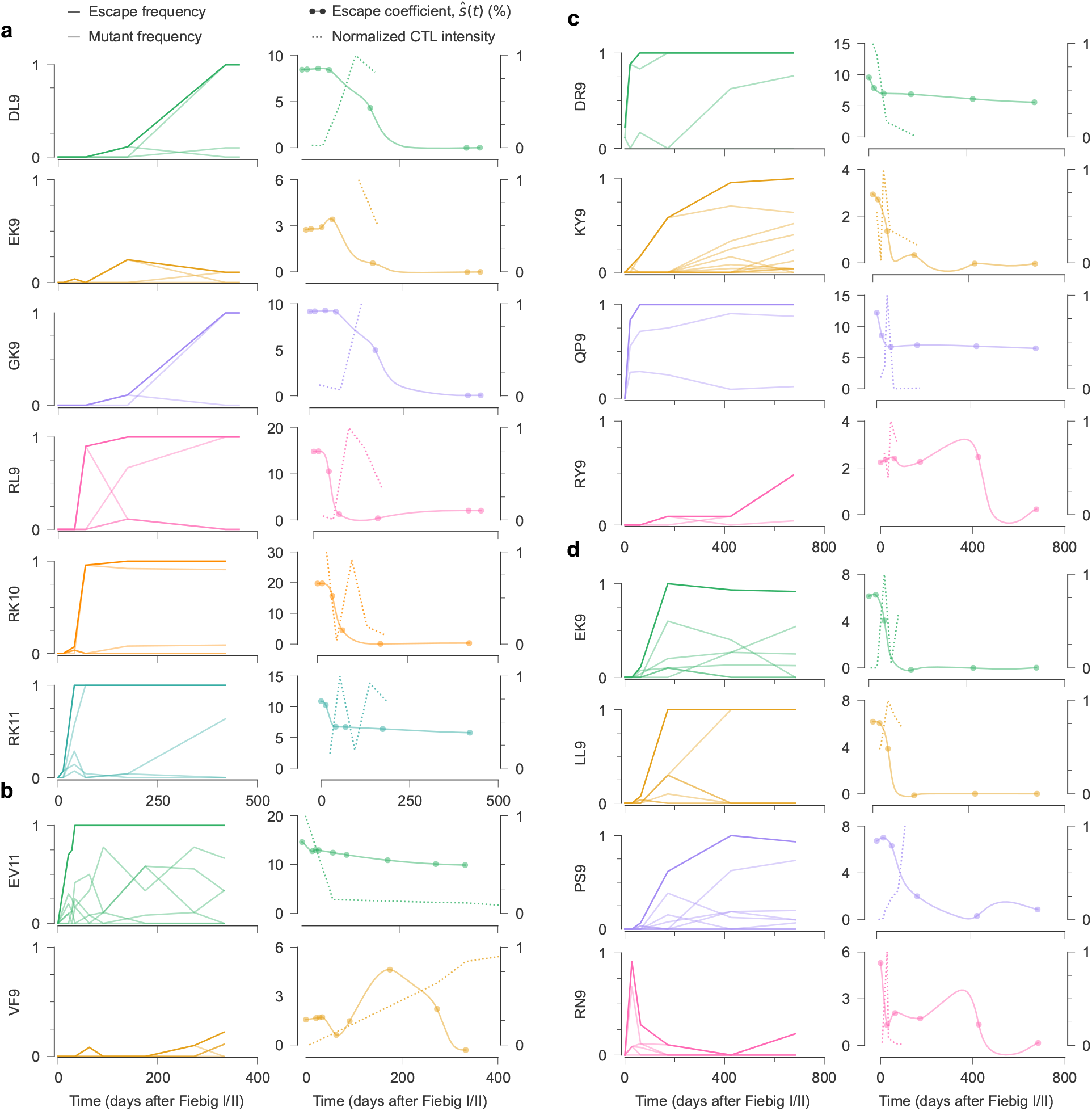
Comparing time-varying selection for CTL escape and experimentally measured CTL intensity. For each CTL epitope with measured intensity data ^60^, we show the frequency of individual escape mutations and the overall fraction of viruses with one or more escape mutations (left column) along with the inferred escape coefficient and normalized CTL intensity over time (right column). This figure shows CTL epitopes for donors (**a**) CH470, (**b**) CH131, (**c**) CH042, and (**d**) CH256.

**Supplementary Fig. S7.**
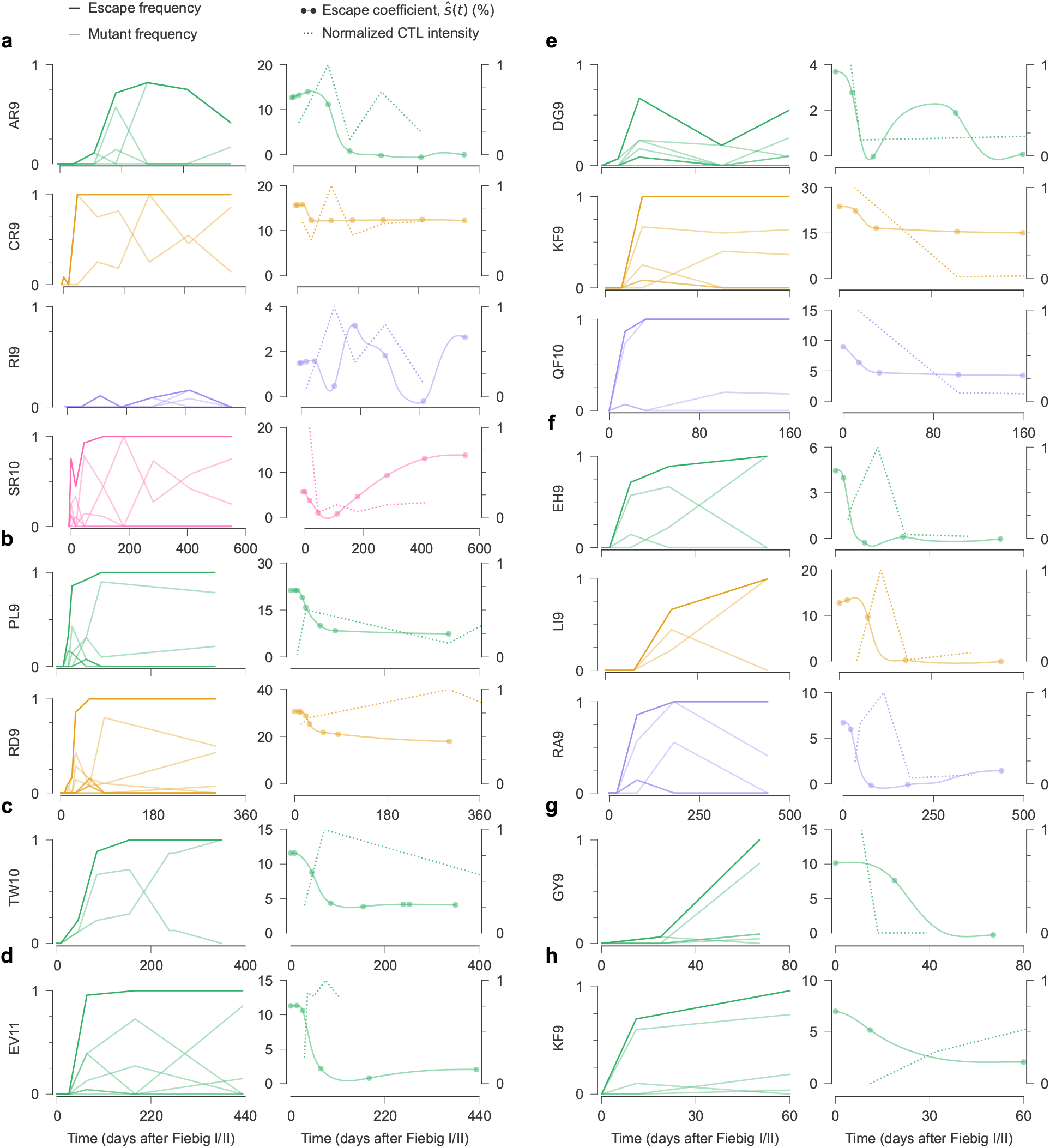
Comparing time-varying selection for CTL escape and experimentally measured CTL intensity. For each CTL epitope with measured intensity data ^60^, we show the frequency of individual escape mutations and the overall fraction of viruses with one or more escape mutations (left column) along with the inferred escape coefficient and normalized CTL intensity over time (right column). This figure shows CTL epitopes for donors (**a**) CH040, (**b**) CH159, (**c**) CH058, (**d**) CH164, (**e**) CH077, (**f**) CH162, (**g**) CH185, and (**h**) CH198.

**Supplementary Fig. S8.**
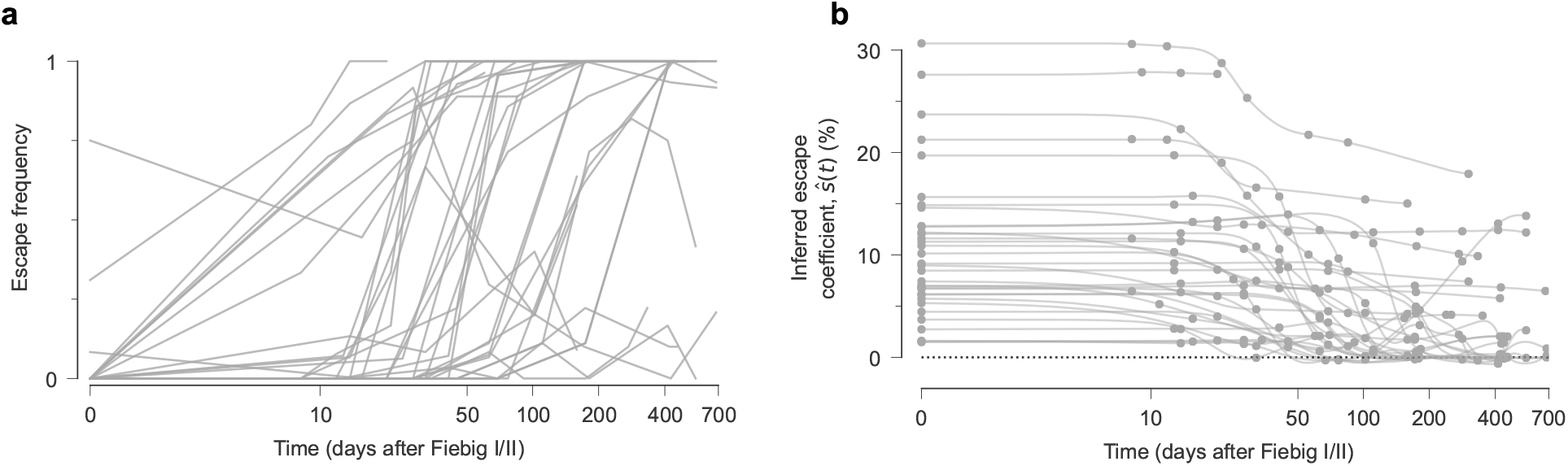
Selection for CTL escape weakens over time. **a**, Frequency trajectories for all CTL epitopes with fitness effects that could be independently estimated from data. **b**, Inferred escape coefficients over time. The pressure for immune escape is nearly always strongest during early infection but exhibits a general trend toward weaker selection over time.

**Supplementary Fig. S9.**
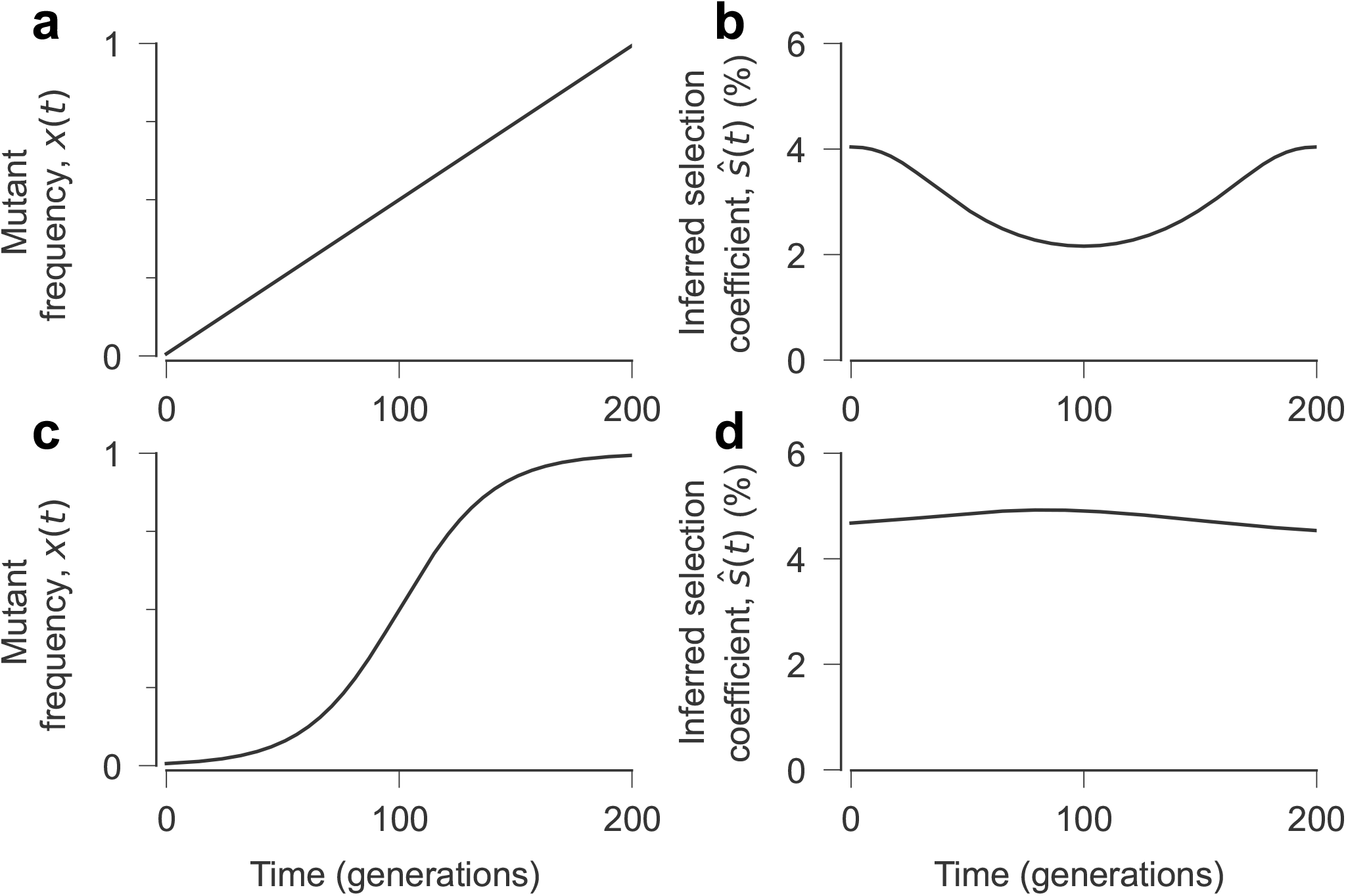
Distortion of inferred selection when allele frequencies are assumed to change by large amounts linearly in time. **a, c**, Mutant allele frequency trajectory. The frequency increases linearly in **a**, while following a sigmoid function in **c. b, d**, The inferred selection coefficients. Both simulations represent single allele cases without mutation or recombination. A linear increase in allele frequency does not result in a constant inferred selection coefficient; however, a sigmoid frequency trajectory yields a nearly constant inferred selection coefficient. The latter result is expected, as the evolution of a single allele frequency with a constant fitness effect should follow a sigmoidal trajectory in the absence of noise and genetic background effects. Compared to **c**, the frequency in case **a** exhibits more rapid changes in frequency during the first and last parts of the trajectory and slower ones in the middle, explaining the non-constant inferred fitness effect in **b**.

